# Binding of single-mutant epidermal growth factor (EGF) ligands alter the stability of the EGF receptor dimer and promote growth signaling

**DOI:** 10.1101/677393

**Authors:** Stefano Pascarelli, Dalmira Merzhakupova, Gen-Ichiro Uechi, Paola Laurino

## Abstract

The Epidermal Growth Factor Receptor (EGFR) is a membrane-anchored tyrosine kinase that is able to selectively respond to multiple extra-cellular stimuli. Previous studies have indicated that the modularity of this system is affected by ligand-induced differences in the stability of the dimerized receptor in a process known as “Biased signaling”. However, this hypothesis has not been explored using single-mutant ligands thus far. Herein, we developed a new approach to identify residues responsible for functional divergence combining the conservation and co-evolution information of ortholog and paralog genes encoding the epidermal growth factor (EGF) ligand. Then, we mutated these residues and assessed the mutants’ effects on the receptor by employing a combination of molecular dynamics (MD) and biochemical techniques. Although the EGF mutants had comparable binding affinities to the wild type ligand for EGFR, the EGF mutants induced a different phosphorylation and cell growth pattern in multiple cell lines. The MD simulations of the EGF mutants show a long-range effect on the receptor dimer interface. For the first time in this study, a single mutation in EGF is shown to be enough to alter the activation of the pathway at the cellular level. These results also support the theory of biased signaling in the tyrosine kinase receptor system and demonstrate a promising new way to study ligand-receptor interactions.

## Introduction

The Epidermal Growth Factor (EGF)-like domain ligand-receptor signaling system is involved in many biological events in multicellular organisms (1). This system is constituted by one receptor (EGFR) and seven distinct peptide ligands (EGF, Epidermal Growth Factor; HBEGF, Heparin-Binding Epidermal Growth Factor; EPGN, Epigen; BTC, Betacellulin; EREG, Epiregulin; AREG, Amphiregulin). Upon binding to the receptor, these ligands can activate multiple intracellular downstream pathways through a network of intramolecular interactions with several feedback loops (2). For all ligands, binding induces a transition in the EGFR from monomer or inactive dimer to an active dimer state (3,4). However, different ligands are able to promote divergent outcomes, even at saturating concentrations; thus, the mechanism responsible of the modular downstream pathway activation is independent of ligand affinity or potency, and likely encompasses intrinsic effects (5). It is well known that the EGFR system plays a key role in cancer development. In particular, some studies have shown that overexpression of EGFR or its ligands may induce different types of cancer (6). A better understanding of the interaction between EGFR and its ligands could lead to the development of targeted therapies (7).

Protein-protein interactions (PPIs), such as that between EGFR and its ligands, are well-studied examples of molecular co-evolution in biological systems. These interactions are sometimes defined by one part (receptor) that binds several counterparts (ligands). In these instances, receptor and ligands experience different selective constraints, where receptors tend to evolve more slowly due to the necessity of binding multiple ligands (8). Furthermore, paralogs, proteins related by a duplication event, are less likely to retain the same function as orthologs, proteins related by a speciation event (9). Then, the paralogous ligands rather than the receptor are a good candidate to test the functional divergence in the biased signaling system. However, this approach is susceptible to indirect factors, such as different ligand binding selectivity, thus resulting in the activation of unrelated pathways. In this work, we decided to test single mutants of one ligand, EGF, by identifying and modifying sites responsible for a divergent function among paralog ligands. Recent studies have shown that some EGF residues like Arg-41 and Leu-47 are highly conserved and important for high binding affinity to EGFR (10). Another study highlighted Tyr-13, Leu-15 and His-16 in EGF as essential for downstream activity of ErbB1 (11). These outcomes were based on structural analyses of ligands and experimental validation. While bioinformatic tools such as contact prediction (12) or molecular dynamics (13) can give a good overall picture of ligand-receptor interactions, the contribution of single ligand residues to the modularity of the system still remains unclear.

Ligand-induced selective activation of downstream pathway has been observed in G-protein coupled receptors, a phenomenon known as “biased signaling” (14,15). A recent report hints that this mechanism might also take place in the EGFR tyrosine kinase (16). Initially, the main contributor to the modularity of the system was thought to be the affinity of the ligands to the receptor. However, the discovery of the ligand EPGN, which induces a potent mitogen effect despite low binding affinity (17), and multiple cell-line studies (5) have changed this perspective. A plausible explanation was initially formulated by Wilson *et al*., where different EGFR ligands induce different receptor dimer stabilities, altering the phosphorylation dynamics (5). Later, new experimental evidence supported this theory by observing a transient activation in stable receptor dimers induced by EGF; whereas the activation of the weak receptor dimer induced by EPGN or EREG persisted for a longer time (18). A difference in dimer stabilities could result in alterations of the receptor oligomerization state, previously shown as a determinant of the intracellular kinase phosphorylation pattern (19). Now, the open question is how these ligands induce different dimer stabilities. Interactions between the interface of the ligand and EGFR have been shown to influence the stability of the dimerized receptor (20), a factor that is related to ligand-specific signaling bias (18,21). It is known that the EGFR dimer can be observed in a symmetrical, “flush” conformation or an asymmetrical, “staggered” conformation (22) depending on the presence/absence of the membrane, glycosylation, and the number of bound ligands (23). Whether and how this plays a role in biased signaling is still unknown.

Here we show how single amino acid substitutions on the ligands effect the biased signaling in the EGF-EGFR ligand-receptor system. We developed a new bioinformatic tool to analyze the co-evolution of the ligand-receptor pair and identify candidate function-altering mutations. The identified mutants induced an altered phosphorylation dynamics and different cellular phenotype. This is the first study to explain differences in biased signaling of EGFR using single-residue EGF mutants. Furthermore, our co-evolutionary analysis can be applied readily to other ligand-receptor interactions.

## Results

### DIRpred

Firstly, we developed a method for predicting residues that are likely to be responsible for functional divergence among paralogs sharing a common interactor (referred as ligands and receptor from now on). We called the method DIRpred (Divergence Inducing Residues prediction). Our approach combines residue-specific conservation measures to identify positions that are conserved among orthologs while diverging among paralogs. The DIRpred analysis is based on the assumption that conservation of a residue in orthologs of a specific ligand shows whether a residue is important for either structural or functional reasons, while conservation of a residue among paralogous ligands denotes the importance of a residue for interaction with the common interactor (the main shared property of all ligands). Thus, residues that are highly conserved in orthologs but not in paralogs of a ligand are likely related to the ligand’s specific function. Unlike other existing methods for prediction of specificity determining residues (a review can be found in (24)), we included inter-protein coevolution measures in order to narrow down those residues that are responsible for a specific interaction. The DIRpred score is calculated as the sum of the four components (I: orthologs conservation, II: complement of the paralog’s conservation, III: ligand-receptor co-evolution, IV: complement of ligand internal co-evolution). Optionally, the analysis can be conducted using a structural alignment of the paralogs (MSTA) instead of the sequence alignment (MSA). An implementation of the DIRpred analysis was done using Python. The pipeline accepts multiple sequence or structure alignments and a reference protein to conduct the analysis. The output consists of a single tabular file containing the four individual scores and the combined prediction score for each protein site, and a plotted recap of the results (Figure S1).

### DIRpred analysis of EGF

We applied the DIRpred algorithm to predict the residues in EGF that induce functional divergence among paralogs (Figure 2). Since the analysis requires prior knowledge of paralogs and orthologs of the target gene, we first performed a phylogenetic analysis to confirm that the reported paralogous ligands of EGF are monophyletic (Figure S2). The same tree was used in the statistical validation of the individual DIRpred scores. A random 53 amino acid long sequence was evolved 100 times on the EGFR ligands phylogenetic tree using Pyvolve (25). Assuming that the distribution of DIRpred scores will be normal, we estimated the probability that a site without functional constraints would have the same or higher than observed score (P-value) (26). We included the results of the statistical analysis to the output of DIRpred for EGF (Figure S3, Table S1). The analysis highlighted residue Asn-32, Asp-46, Lys-48 and Trp-50 as potential candidates for paralog functional divergence. Asn-32 and Lys-48 show a very small conservation in both sequence and structure alignments resulting in a high partial score (II). Asp-46 has a relatively high coevolution with the receptor (III), while Trp-50 has high scores overall (Figure S4).

**Figure 1.**
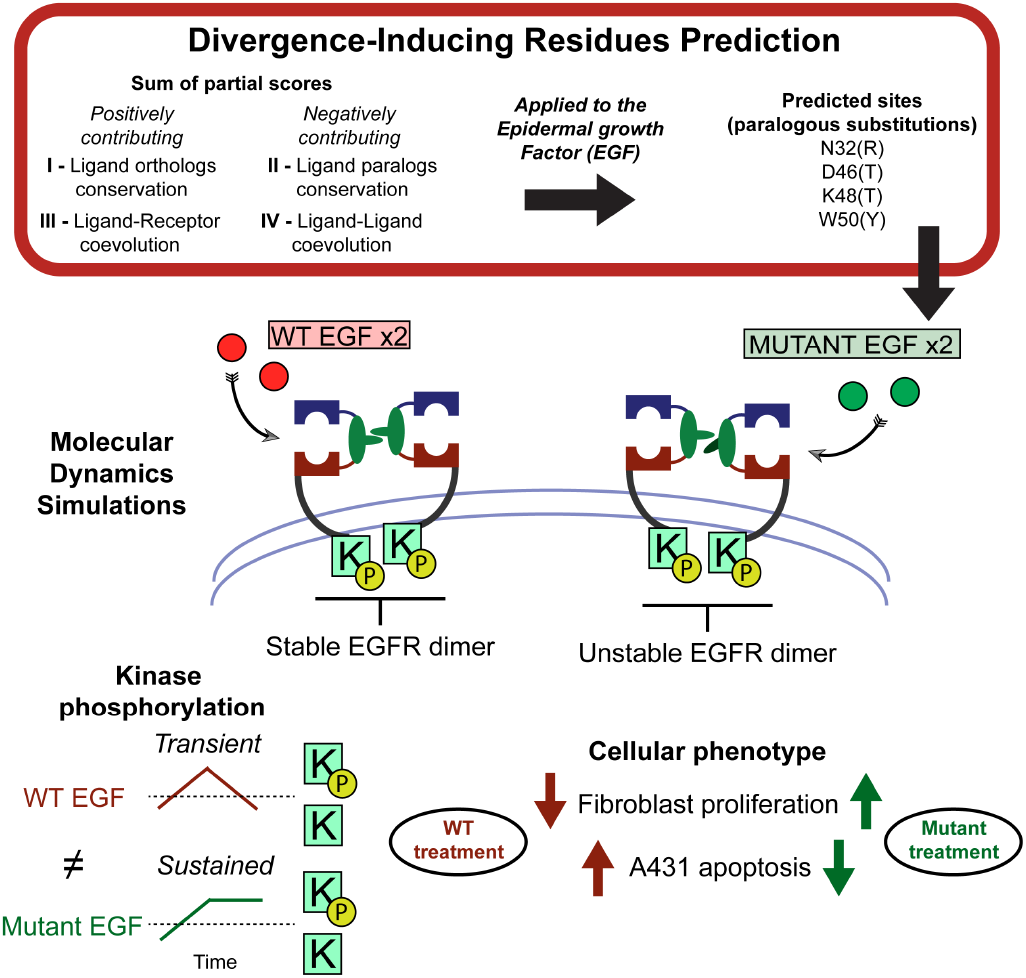
Study rationale. EGF mutants were identified using an *ad hoc* methodology that combines conservation and coevolution measures, DIRpred. Unlike WT EGF, the mutant ligands induce an unstable conformation of the receptor dimer, similarly to a dimer with a single WT EGF. Treating A431 and fibroblast cells with either WT EGF or mutant EGF resulted in a different response both in the phosphorylation dynamics and the cell proliferation phenotype.

**Figure 2.**
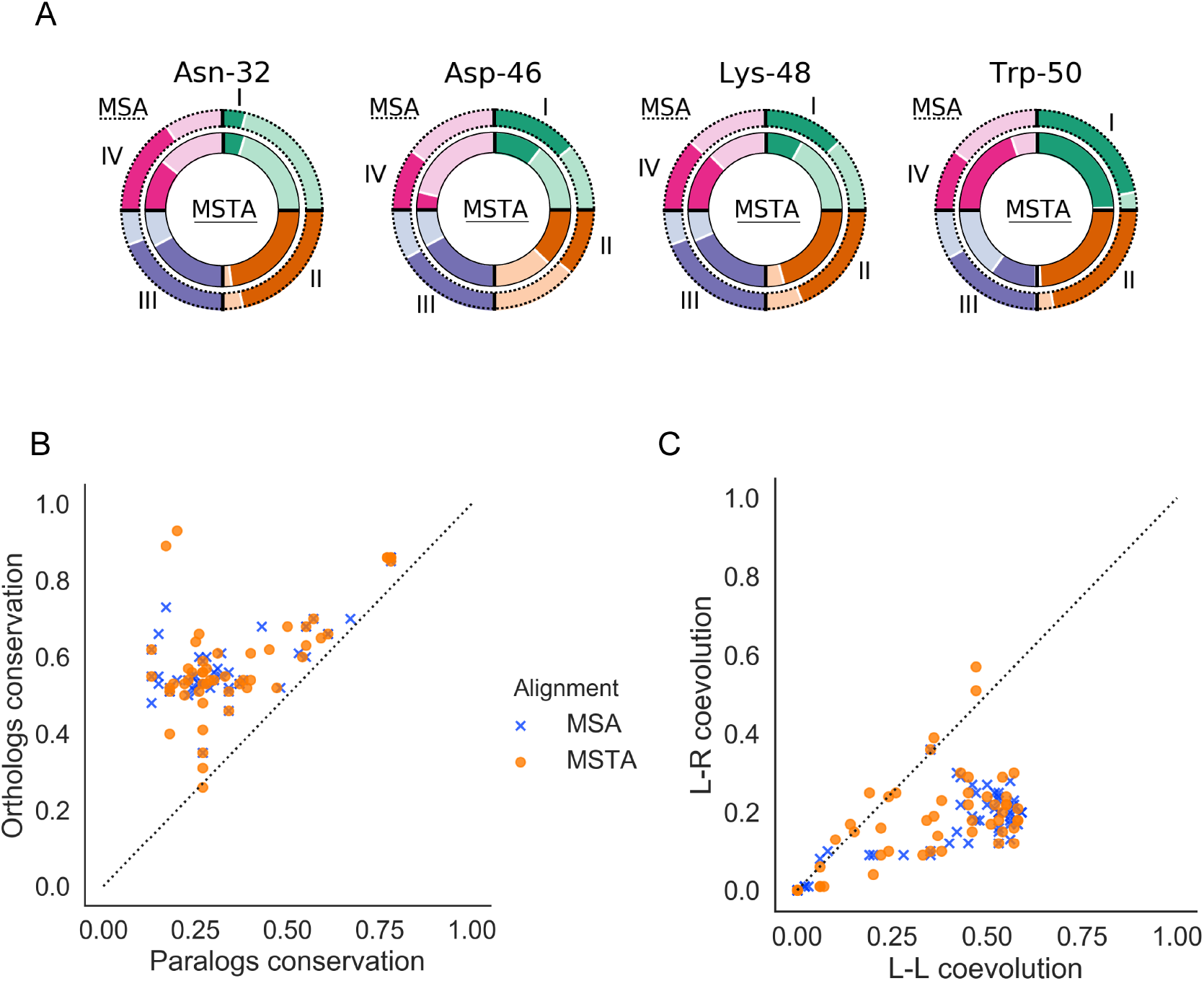
DIRpred analysis of EGF. (A) The relative ranking score of the four mutants. The outer circle represents paralogs multiple sequence analysis (MSA)-based scores, while the inner circle represents paralogs multiple structural alignment (MSTA)-based scores. The darker color indicates a higher ranking in the EGF sites-based scoring. The ranking values are reported in Table S2. (B) Cross-conservation plot. The plot is obtained by crossing the two conservation scores. Interestingly, no point lies in the bottom right half of the plot (high paralogs conservation and low orthologs conservation), suggesting that paralogs and orthologs conservation are not independent. This observation points out that there is no organism-specific adaptation shared by all ligands at the protein sequence level. (C) Cross-co-evolution score. The plot is obtained by crossing the two co-evolution scores, the ligand-receptor (L-R) co-evolution score (III) on the y axis and the ligand-ligand (L-L) co-evolution score (IV) on the x axis.

Three out of the four positions were mentioned in previous reports, however none of them was individually mutated. The tryptophan in position 50 is a strong outlier in our bioinformatics analysis, along with Trp-49 (Figure 2). Their score is high even when using conservation measures that do not take amino acid type change into account (data not shown). However, Trp-50 was a better candidate for testing because of its outward facing position, while Trp-49 is involved in buried protein contacts (27). Trp-49 and Trp-50 could be responsible in facilitating the interaction of EGF with the membrane phospholipids, as it happens for Pro-7 and Leu-8 (28); Trp-49 and Trp-50 are not burying inside the bilayer when EGF and a membrane are in solution alone (29), though this might be different when in complex with the receptor. For example, this could be achieved by Trp-50 through the formation of a helix, clustering together with Val-34 and Arg-45 around the conserved Leu-47 (30). Mutation N32R is on the interface between ligand and receptor. The slightly higher affinity is probably due to the presence of the guanidinium group of R which is positively charged and could interact with Gln-16 of EGFR ECD (Figure S5). Interestingly, mutations of the corresponding position in chicken TGFA were able to alter the mitogenicity without strongly affecting the binding affinity (31). While no previous literature reported about Asp-46 before, Lys-48 was found in two mutants that showed higher affinity (32).

Next, we manually chose which amino acid to introduce on the four sites depending on several factors. The main contribution to the decision was given by the paralogs alignment, while also considering the amino acid type and the ligand functional divergence. The paralogs were divided into two groups based on their kinetics parameters of interaction with the receptor. After that, we selected an amino acid that infers a significant change in biochemical properties and that is found in the paralogs group without EGF. When multiple choices were possible, priority was given to EPGN or EREG, since these two ligands are the ones observed to induce a biased signaling in Freed *et al* (18). The four designed EGF mutants with a single amino-acid substitution that were selected for functional characterization are N32R, D46T, K48T and W50Y.

### Biochemical properties of the EGF mutants

To determine the integrity of the secondary structure of the mutated ligands and the functional effects of these amino acid substitutions, we first performed *in-vitro* analysis. Initially, Circular Dichroism (CD) spectroscopy was used to confirm that the secondary structure of the mutants was maintained (Figure S6). The β-sheet content of the EGF mutants (ranging from 0.41 to 0.54 %) was not substantially different than the WT EGF (0.44 %) while the other secondary structure varies. It is not surprising that only the beta sheet content can be detected by CD considering that EGF is a peptide (53 amino acids) constituted by two β-sheets connected by a loop and the regions at the C- and N-terminus are flanking. Then, we tested the ability of each mutant to bind the soluble extracellular domain of EGFR, by Isothermal Titration Calorimetry (ITC) and MicroScale Thermophoresis (MST). The mutants bound the receptor analogously to the WT sample in both the experiments (Figure 3). Even though the N32R sample showed a slightly steeper response in both the experiments, the mutations did not appear to strongly affect the ligand secondary structure and the ability to bind the receptor.

**Figure 3.**
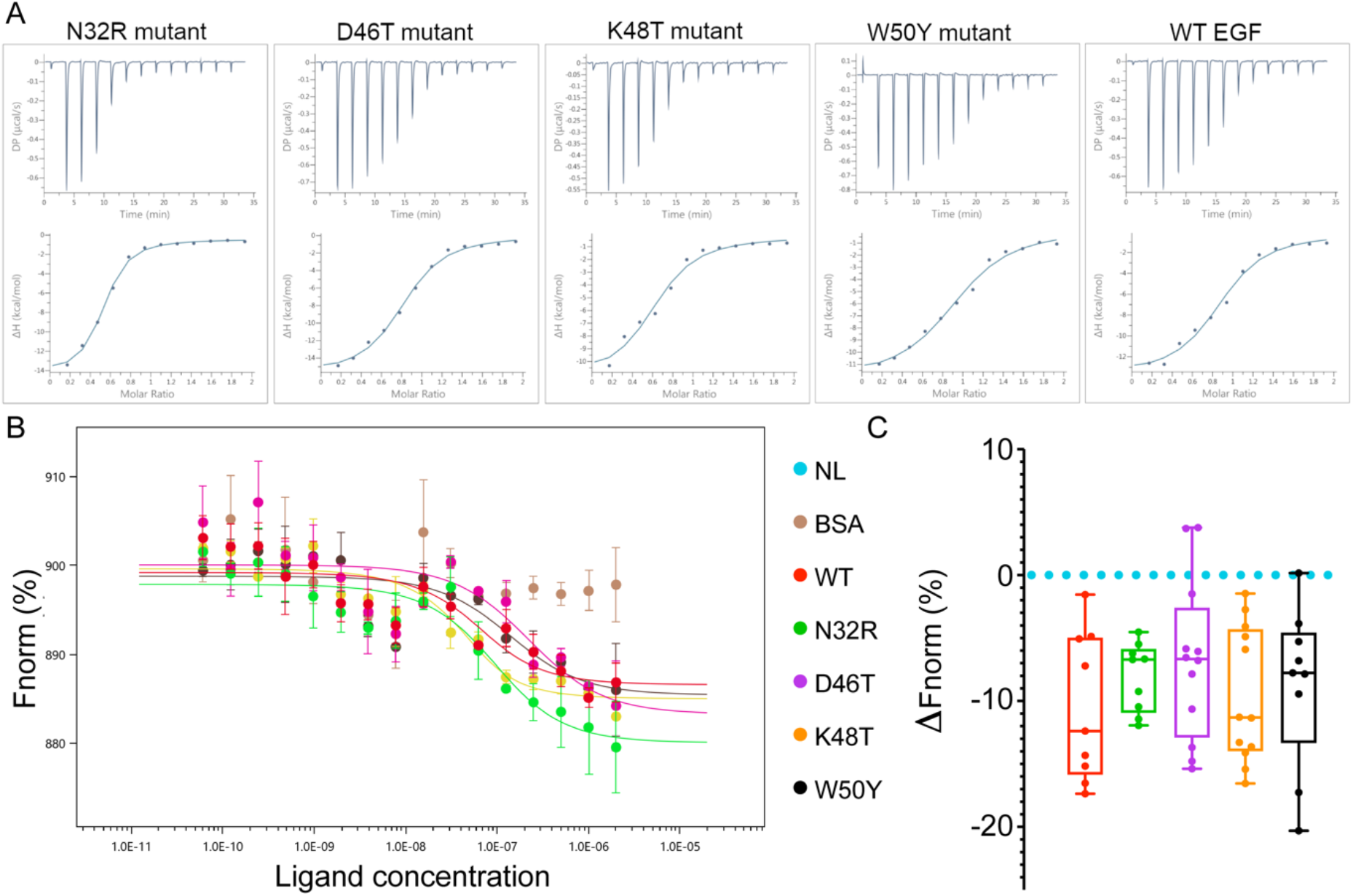
Binding measurements of EGF mutants to the EGFR receptor by Isothermal Titration Calorimetry (ITC) and MicroScale Thermophoresis (MST). (A) ITC analysis of WT (Wild Type) EGF ligand, and mutants N32R, D46T, K48T, and W50Y binding to the ECD of the EGFR receptor at 25° C. Measurements were taken by adding WT EGF or mutants at 200 μM to the ECD of EGFR at 20 μM. (B) Extrapolated curves of the MST experiment. The normalized fluorescence difference (Fnorm) at 20 seconds for different concentrations of ligands was analyzed using the NanoTemper Technologies analysis software. Using the Kd model, it was possible to fit a curve for every sample except BSA, that showed no binding. All ligands show a transition at about 100nM. (C) Multiple sampling of thermophoresis was performed at a concentration of 100nM, the point of the curve where we expected to observe a biggest difference for a ligand with an altered affinity. The Fnorm is shown in relationship to a NL (No Ligand) sample average coming from the same experimental batch.

### Biological effects of the EGF mutants

To shed light on the biological outcome of the short-term response, we sampled the effect of the EGF single mutants on EGFR expression and phosphorylation in A431 cells. A431 is a human epidermoid carcinoma cell line that overexpresses EGFR. For this reason, we postulated that any changes at the receptor level would be amplified. We measured the amount of total and phosphorylated EGFR protein at multiple timesteps by Western Blot (WB) after treating with saturating concentrations (100 nM) of WT or mutant ligands (Figure 4). Remarkably, D46T mutant showed a reduced level of EGFR phosphorylation (pEGFR) up to 30 minutes after treatment (Figure 4A). Meanwhile, the pEGFR bands for K48T and W50Y samples appear slightly stronger than the WT in the first three timesteps. However, the differences are not significant. The dimerization state of the receptor, tested by cross-linking assay, follows a similar trend as the phosphorylation experiments (Figure 4B). After 1 h or 6 h, the treatment with the ligands also caused a reduction in the total amount of EGFR compared to the control (Figure S7). However, the reduction of D46T sample did not appear as marked as the other samples. This result shows an inverse relationship between receptor activation and receptor quantity, a fact that could be explained by the biased signaling theory. Thus, while the EGF mutants showed comparable binding affinities to the receptor, at least one of the mutants showed a different phosphorylation pattern compared to the WT ligand. Furthermore, the observed differences in the total amount of EGFR protein indicate a possible alteration in the membrane expression or recycling of the receptor.

**Figure 4.**
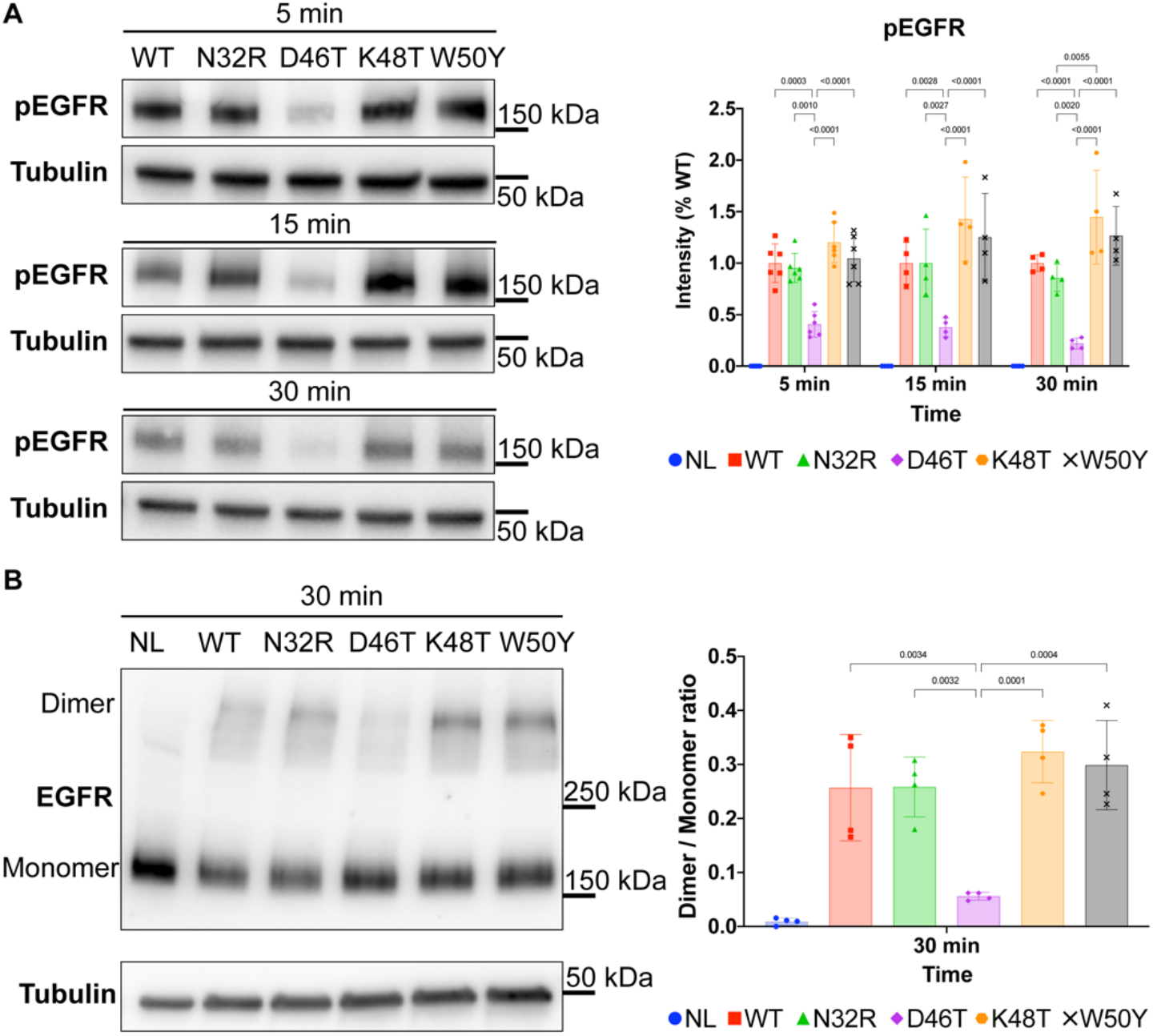
Short-term response: Effects of WT and single-mutant EGF on the receptor phosphorylation and dimerization in A431 cells at different time points. (A) Short-term effects on the phosphorylation level of EGFR Tyr-1173 after treating A431 cells with 100nM concentration of different ligands. The membrane containing EGFR and tubulin was separated after the transfer. D46T treated cells show a statistically significant reduction in the level of phosphorylation compared to the other samples. (B) A431 cells were treated with 100nM WT or mutant ligand. After 30 minutes, the cell lysate was run through a protocol for cross-linking and western blot. The bars represent mean ± s.d. of at least four biological repeats. The number on top of the bars shows the p-values of a 2-way ANOVA (A) or one-way ANOVA (B) multiple comparison corrected for multiple sampling using the Bonferroni correction. Details of the ANOVA statistics can be found in Table S3. Band intensity estimates were calculated using BIO-RAD ImageLab software (BioRad). Plots and statistics were performed using PRISM software (GraphPad).

In order to observe the long-term effects of the mutants on the cellular phenotype, we performed a cell growth assay using an IncuCyte^®^ live-cell analysis system on two fibroblast cell lines (Bj5-tα and Albino Swiss mouse 3T3). With the same platform, we also performed an apoptotic assay on A431 cells by measuring the reduction in cell population, using the Annexin V green reagent to confirm the induction of apoptosis. After one day of initial incubation, we subjected the cells to 1, 10 or 100 nM concentration of growth factors. Treating Bj5-tα fibroblasts with 100 nM WT EGF resulted in the highest reduction in the proliferation rate compared to the control. The reduction in proliferation was accompanied by a change in cellular morphology that could be a signal of differentiation (Supplementary Data 1). However, treatment with any of the mutants decreased the ability of the ligand to suppress growth (Figure 5A). The D46T and K48T mutants differed most from the WT (Figure S8A). We observed a similar trend in the Albino Swiss mouse 3T3 cell line (Figure S8B). In both cell lines, the effects are concentration dependent (Figure S9). Next, we further tested the mutants in an apoptosis assay using the A431 cell lines. This cell line is known to exhibit apoptosis when treated with high (>10 nM) concentrations of the EGF peptide (33). The WT showed the highest decrease in cell population 48h after treating with 100 nM WT, while the four mutants had an intermediate response between WT and the control. Cells treated with the W50Y mutant have the closest level to the WT (Figure 5B). In WT EGF cells, we observed evidence of apoptosis in cell’s globular processes and increased ratio of Annexin to confluence signal (Figure S10). Meanwhile, D46T and K48T treated cells displayed signs of cellular differentiation (Figure 5C), in comparison to the no ligand and other mutants (Figure S11, Supplementary Data 1). Significantly, these results show that single amino acid changes in the EGF ligand display differential effects on the EGFR transduction mechanism.

**Figure 5.**
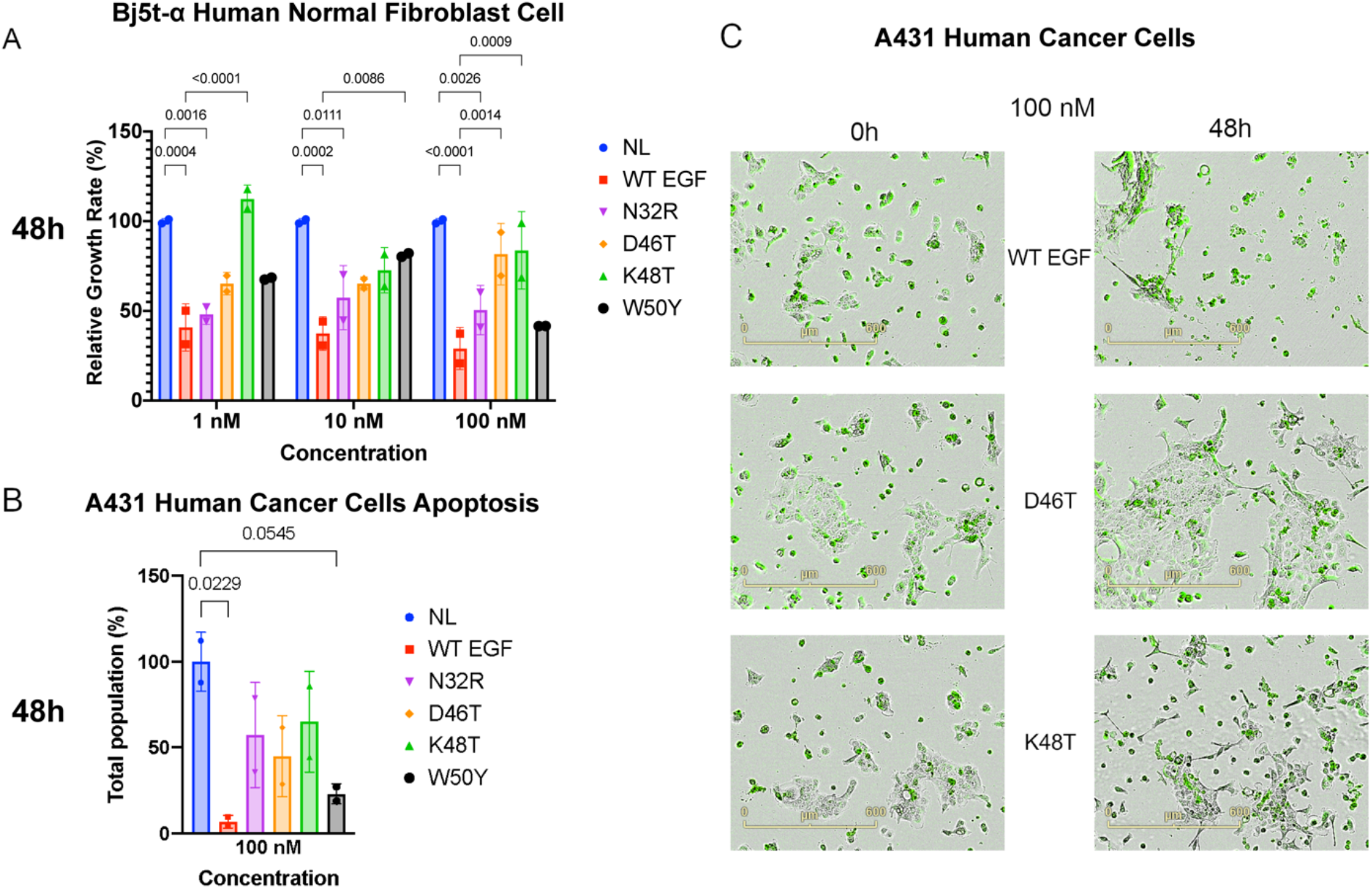
Long-term response: Growth assays of cells treated with EGF variants. (A) Effect of different concentrations of EGF variants on proliferation of the human normal fibroblast Bj5-tα cell line. Data represent the percentage of the growth rate, calculated from the confluence of cells, relative to the control (mean ± standard deviation). Percent confluence was estimated 24 h after the treatment (two replicates per treatment). (B) Apoptosis effect of WT or mutant EGF on A431 cells. EGF-induced apoptosis is measured as the reduction in total population compared to the control. (C) Comparison of A431 cell growth after treatment with 100 nM WT EGF and EGF variants D46T and K48T. Cells were labeled with fluorescent Annexin V Green Reagent. Plates were pre-warmed prior to data acquisition to avoid condensation and expansion of the plate, which affect autofocus. Images were captured every 2 h (4x magnification) for 3 days in the IncuCyte system. The number on top of the bars shows the p-values of a 2-way ANOVA (A) or one-way ANOVA (B) multiple comparison to the control lane, corrected for multiple sampling using the Bonferroni correction. Details of the ANOVA statistics can be found in Table S3. Band intensity estimates were calculated using BIO-RAD ImageLab software (BioRad). Plots and statistics were performed using PRISM software (GraphPad).

The observation that the four mutations in EGF alter signal transduction without disrupting any contacts between the ligand and the receptor can be explained by the “loss of symmetry” model of EGFR signaling (18). Freed *et al*. observed how stable symmetrical dimers show a short-lived nature of signaling, while asymmetric dimers conduct a sustained activation, lasting longer than 24 h. Such a model provides an explanation on how a low level of phosphorylation after EGF treatment causes modifications in the apoptotic behavior observed in the cellular growth experiments. Compared with WT EGF, the D46T variant show a constantly lower phosphorylation signal, possibly through the formation of a less stable, asymmetric dimer. Thus, the observed effects of the EGF mutations on EGFR phosphorylation and dimer stability are consistent with the “loss of symmetry” theory. However, the fact that some mutants have a similar phosphorylation pattern as the WT EGF but show different cellular phenotypes suggests that there might be other factors at play.

### Molecular dynamics

To understand how EGF single mutants affect the receptor signaling transduction, we performed full atomistic molecular dynamic (MD) simulations of the extracellular EGFR in complex with WT or mutant ligands (100 ns each). We modeled the receptor starting from the asymmetric, “unstable” conformation of 5WB7 (22). In this way, we expected to observe a fast rearrangement for those simulations where a ligand more favorably induces a stable dimer. In previous literature, an unstable conformation was observed when removing one of the EGF ligands (23). For this reason, we also performed one simulation of the EGFR dimer in complex with only one WT ligand as a comparison (1ligEGF). The MD simulations quickly converged to a stable RMSD (Figure 6A). For each simulation, we calculated the number of H-bonds between the two receptor dimers, and between ligand and receptor. In the WT EGF simulation, the EGFR dimer had an average of 15 H-bonds, higher than any other simulation (Figure 6D). However, the number of bonds between receptor and ligand was not altered (Figure 6E). In addition, during the course of our simulations we also noted differences in the RMSF specifically located at the dimerization arm domain (Figure 6B). The conformational space sampled by the dimerization arm of K48T simulation was much wider than the WT simulation (Figure 6C). To analyze the temporal distribution of these motion, we measured the distance between Pro-272 and Gly-288 of different EGFR monomers. This distance was chosen because it was able to discriminate between the EGFR dimer in complex with EREG (PDB ID 5WB7) and EGF (PDB ID 1IVO). In 5WB7, one of the two distances is much bigger (~ 1.10 nm vs 0.4 nm) compared to 1IVO (Figure 6F). In our simulations, we observe WT showing a sharp peak at 1 nm, while the mutants and 1ligEGF have a secondary peak at higher distances (Figure 6G). Thus, the mutations in EGF appear to affect the stability of the EGFR dimer without affecting the stability of the EGF-EGFR interaction.

**Figure 6.**
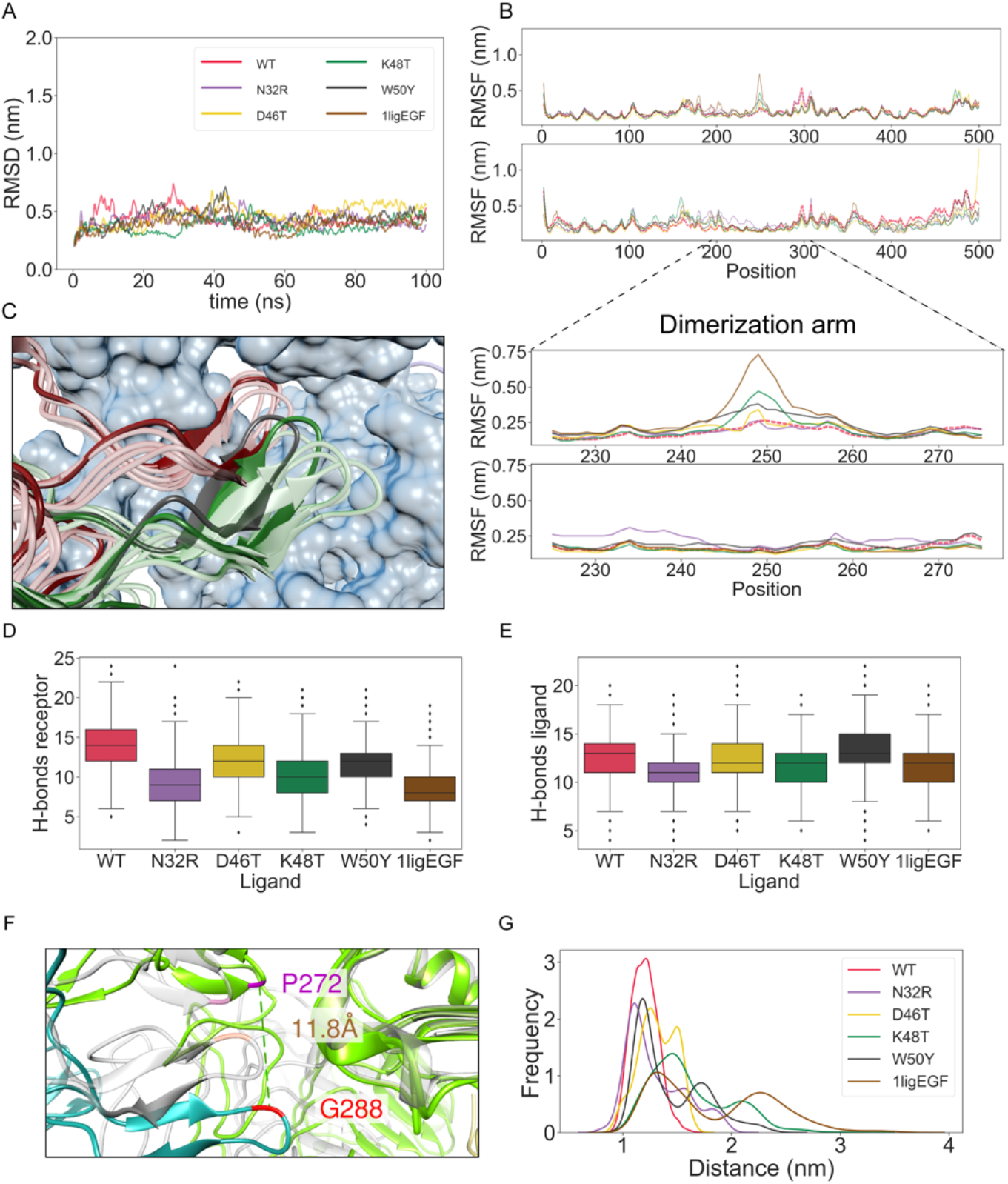
Molecular Dynamics of the EGFR extracellular domain. (A) Root Mean Square Deviation from the initial structure. (B) Top, Root Mean Square Fluctuations of the 100 ns simulations. The WT level is marked in bold and dashed red line. Bottom, focus on the RMSF of the dimerization arm domain. The two plots represent the two EGFR in the receptor dimer. (C) Dynamics of the dimerization arm domain in WT (red) and K48T (green) simulations. A structure every 20 ns of simulation was taken and aligned to 1IVO reference structure (blue surface and black ribbon). The structure at 20 ns is represented in solid colors. K48T simulation shows a more dynamic dimerization arm. (D) Number of H-bonds formed between the two receptors during the 100 ns simulation time. (E) Number of H-bonds formed between the receptor that shows a dynamic dimerization arm and the corresponding ligand during the 100 ns simulation time. (F) Distance used to investigate the fluctuations of the dimerization domain highlighted on the structure. Pro-272 (purple) and Gly-288 (red) are highlighted both on 5WB7 (green and teal) and in 1IVO (grey). The distance between the alpha carbons of the 5WB7 couple is shown in brown and is notably longer than 1IVO. (G) Distribution of the distance between Pro-272 and Gly-288 in all the simulations. WT simulation shows a single peak at ~1nm, while in all other ligands the distance has two peaks.

## Discussion

The prediction of functional residues is a well-developed field (34), where conservation measures are considered a key factor to rely on. Tools such as ConSurf (35) and the ET-like methods (36) are able to identify slowly evolving positions that are involved in folding, interaction, or catalytic activity (34). However, the specific reason why a residue is conserved often remains unclear. In this work, we show a new method to identify residues that affects specific functions in a system of interest. Our approach combines a conservation score calculated from the structural alignment of paralogs and among orthologs, with an intra- and inter-molecular co-evolution score with previously known interactors. Conservation and coevolution give a complementary signal, thus improving the overall predictive performance (37). The coevolution score was introduced to DIRpred to highlight residues that are directly responsible of an interaction with the receptor, at the expenses of those that interact intramolecularly within the ligand. While the first part efficiently identifies Asn-32 and Lys-48 as putative interactors, the second part does not properly give a penalty to residue Tyr-29. The interaction of this position with His-10 is also conserved in EGF but with reversed positioning (Figure S12), resulting in a low paralog conservation (therefore high contribution to DIRpred score). While it is still possible to optimize the co-evolution scoring function, integrating conservation and coevolution measures is a promising way to recall information of specific functions involving protein-protein interactions.

The herein studied mutations do not alter significantly the ability to bind EGFR. However, the mutants showed a different cellular effect on Bj-5ta cells (Figure 5). A delayed proliferation response compared to control might seem counterintuitive for fibroblast (38). Though, Bj-5ta hTERT immortalized fibroblasts have features that distinguish them from *in vivo* fibroblast, like differential gene expression and epiregulin-dependent proliferation (39). A decrease in cell proliferation after EGF treatment could be the result of competition between the endogenously synthesized epiregulin and EGF, thus altering the balance between proliferation and differentiation.

A431 cells constitutionally express EGFR at high levels. Treatment with EGF has been observed to promote STAT1 dependent apoptosis (40). This pathway is dependent on the internalization of the ligand-receptor complex by the endocytosis process, a key factor in the ligand-induced biased signaling (16). In our experiments, A431 cells treated with mutant EGF ligands show a higher growth rate, therefore a decreased rate of apoptosis compared to WT. Given the setting of the experiment, modularity in the endocytosis pathway is a straightforward explanation to the observed differences in growth rate and in protein levels and phosphorylation. Interestingly, D46T and K48T treated cells showed an increase in long and thin cellular processes in all replicates (Figure 5). D46T was also observed with a marked difference in the EGFR phosphorylation compared to WT (Figure 4). A possible explanation could be the EGF-induced tubular formation, an alternative EGF-induced pathway reported in intestinal epithelial cells (41). In both the cellular growth experiments, the growth rate of the mutants is intermediate between the control and EGF, and concentration dependent. This phenomenon, and the cell-line specific response of growth hormones is consistent with previous literature reports (42).

The analysis of the fluctuations in the dimerization arm reveals an underlying bias in ligand-induced receptor dimerization, originally not visible from the static images of structure comparison. K48T is the mutant that induce the biggest deviation from the WT. Although, all mutants show a transition. While giving an initial outlook on the effect of the mutants, our MD simulations do not take into account other factors that could be important in the mechanism of action of EGFR. Differential multimerization (43), oligomerization (19), receptor glycosylation, and the interaction with the membrane (23) are factors where the underlying bias induced by the mutant ligands could also have an effect in.

In this work, we showed how a single mutation of EGF is able to alter the specific functional relationship with the receptor. For functional divergence to arise, it could take as little as mutating 15% of the sequence (44). However, the EGFR ligands divergence date back to the vertebrate ancestor of R1/R2 whole genome duplication, up to 500 million years ago (45), as hinted by the low (~25%) sequence identity. From our results, the sequence distance does not reflect the distance in function. In fact, the functional divergence of EGF was altered with just a single targeted mutation.

In conclusion our data suggests that a single mutant ligand induces a conformational change of the receptor that then affects receptor dimer stability, with plausible effects on phosphorylation level and downstream pathway activation (46). This shows how the persistence of biased signaling in EGFR is in an unstable equilibrium, where the observed conservation of diverging sites among paralogs is naturally reinforced to maintain the functional divergence. To determine whether a short distance in function space among paralogs is a consequence or a necessity for living systems, further studies will be required.

## Experimental procedures

### Sequence and structure analysis

The sequences of EGFR ligands and the multiple sequence alignment of EGF orthologs were obtained from the Ensembl database (47). Multiple sequence alignment (MSA) of all ligands was performed with MAFFT software (48). X-ray structures were obtained from the PDB database (49). Structural alignments were created using Chimera (50).

### Phylogenetic analysis

From the multiple sequence alignment of EGF from different species, nearly identical sequences were removed. The *Drosophila melanogaster* EGF sequence was added as an outgroup in the EGF phylogeny, while *Caenorhabditis elegans* EGF was used as outgroup for the ligand phylogeny. MSA and phylogenetic tree images were created using Unipro UGENE software (51). Three phylogenetic trees were made using the Neighbor Joining (NJ), Maximum Likelihood (ML), and Bayesian (MrB) methods in IQTREE (52), using ModelFinder to scan for the best-fit evolutionary model and parameters (53).

### Divergence Inducing Residue PREDiction

We identified sites in EGF responsible for functional divergence using a method that combines evolutionary and co-evolutionary data, called DIRpred (Divergence Inducing Residue PREDiction). The DIRpred scoring function combines four components to evaluate each residue: I) The combined conservation scores in the ortholog alignments. This score is calculated by averaging the conservation of a reference ligand site with the conservation in the respective positions of the other ligands’ orthologs alignments; II) The complementary value (1 – x) of the conservation score in the paralogs alignment. This score can be obtained either from sequence alignment (paralogs MSA) or structural alignment (paralogs MSTA). III) The highest co-evolution score between a reference ligand site and all of the receptor sites. IV) The complementary value (1 – x) of the highest co-evolution score between a reference ligand site and all of the other ligand sites in the joint orthologs alignment of all ligands (Figure 7).

**Figure 7.**
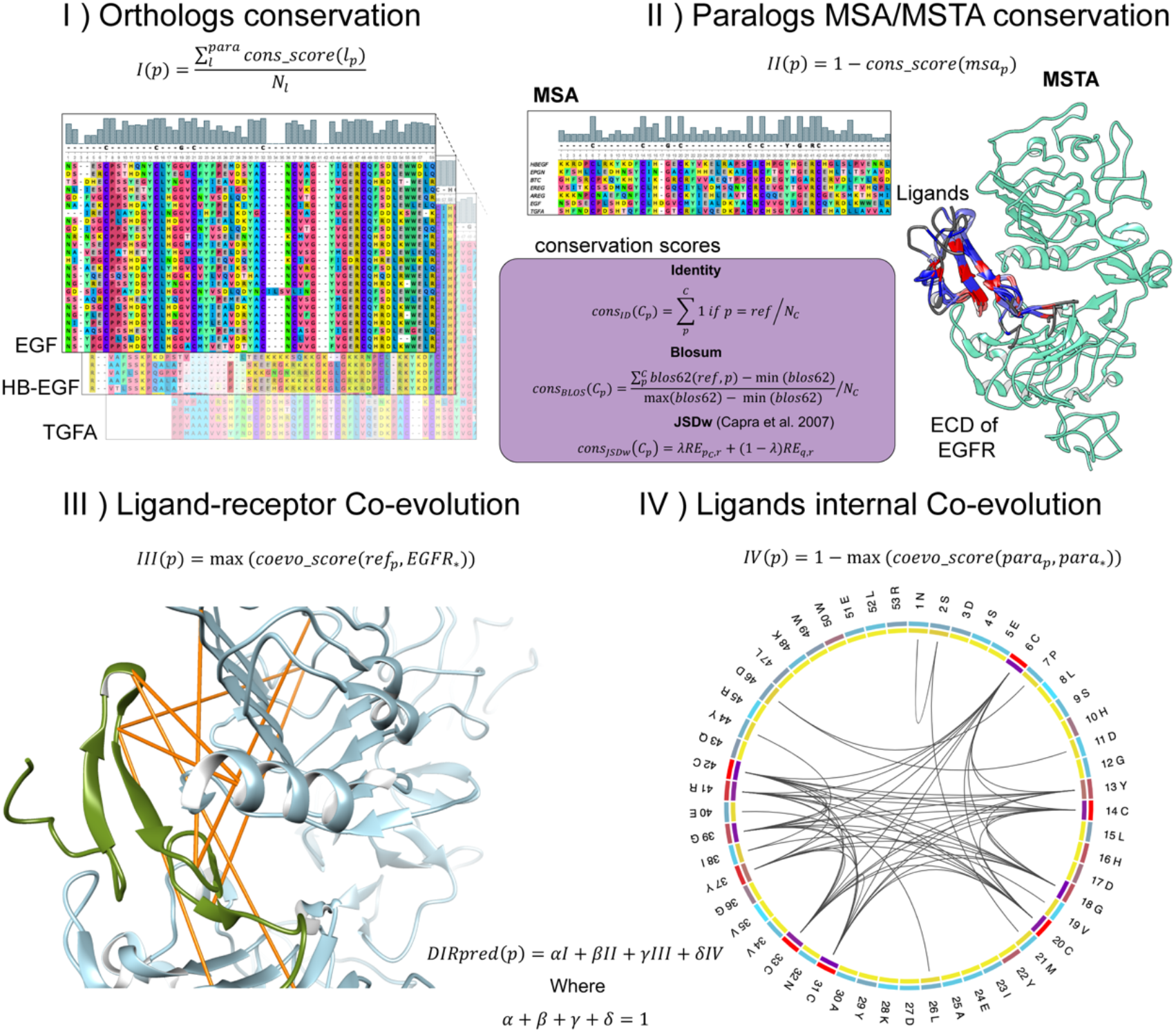
DIRpred rationale. The prediction of Divergence Inducing Residues consists of a linear combination of four site-specific scores, in roman numerals. In I, the orthologs MSA of each paralog is used to compute a conservation score (as exemplified from the histograms on the alignments). Alignment images were produced using Unipro software (72). In II, the human paralog sequences or structures were aligned and the same conservation score function was used. In the purple box, the conservation score functions are represented. In III, the co-evolution score between EGF and each receptor site was computed using Mutual Information (MI). The yellow lines on the structure connect the highest scoring co-evolving residues between ligand and receptor. In IV, MI co-evolution score was computed from the combined ligands orthologs alignment. The circular plot was performed using MISTIC2 (73).

Conservation scores were calculated using three different formulations: 1) IDENTITY, 2) BLOSUM, and 3) JSDw. The IDENTITY score measures the frequency of appearance of a reference residue in the MSA. The BLOSUM score takes the amino acid substitution frequency into account using the BLOSUM62 matrix, normalized by the maximum and minimum score for the BLOSUM matrix. The JSDw score is based on Jensen Shannon divergence with a window of residues (54). The fasta sequences were imported using BioPython package, while analysis and plots were performed with Python3 package SEABORN. EGF index positions from the paralogs MSA and MSTA were respectively used to align the MSA and MSTA based scores. A schematic representation of how the four scores were obtained is showed in Figure S1.

The code used in the analysis of the DIRpred score and plots and the data used in this paper are shared on Github: https://github.com/oist/DIRpred.

### Statistical validation of EGF DIRpred

The statistical analysis of EGF DIRpred scores was done as in Mirny and Gelfand (55) with some modifications. The key point is that a null-hypothesis dataset should take into account the higher similarity found between orthologs, rather than between paralogs. Such a dataset was obtained through a simulated evolution performed using the Python package Pyvolve (25). The sequences in the output dataset were required to have a relationship akin to those of the EGF homologs. Using the WAG model (56), a random 53 amino acid-long protein sequence was evolved from the root of the EGF homologs tree until each leaf node, simulating a scenario of neutral evolution. Then, the sequences were divided in orthologs and paralogs using the original tree classification. The output was used as input for the DIRpred pipeline to obtain four partial scores for each site. After 100 times repetition, the partial scores were gathered together to form four background distributions. Through the assumption of normality, it was possible to estimate the probability of a site to have a higher or equal score as a randomly evolved site, i.e., without functional constrains (26). The scores were considered significant only on the side where they contribute positively to the total DIRpred score, for example, when there is a lower level of paralogs conservation than expected. For this reason, only one side of the normal distribution is used for the calculation of the P-value. (Figure S3). In this way, it was possible to isolate the highly conserved cysteine positions, that usually get a misleading high overall DIRpred score. About the relative contributions of individual scores, an equal weighting system was preferred over the use of arbitrary values, which might not be always optimal. However, the pipeline allows the user to provide his own weighting system that might be more optimized for the user’s study subject.

### Selection of the mutations

Along with the DIRpred score, the choice of positions for mutation was influenced by two manually curated factors: the distance from the receptor and the amino acid variation among ligand types. We designed mutations with the aim of inferring a transition to the amino acid properties in sites where the ligands show a different pattern. Overlapping residues at a given position were divided into two groups, based on an EGF-like and non EGF-like stabilization of the receptor dimer. This property was previously shown to follow binding affinity (18). Residues that introduced a noticeable shift in amino acid properties between the two groups were selected. For example, position N32 is hydrophobic in the high affinity ligands group, positively charged in the low affinity group and negatively charged in EGF. Finally, we carefully analyzed exceptional cases in the DIRpred scoring. Some of the residues that show high score have intramolecular interactions with another amino acid in the ligand. These residues, if mutated, will lose EGF structural stability (namely “residue swapping” behavior showed in Figure S12). The decision of which mutation to introduce was made using the paralogs alignment, with a preferential choice over the residue found in EREG or EPGN. Positions 32, 48 and 50 have high DIRpred score. Position 46 was included although having a lower score, because the substitution pattern matches the two ligands groupings. Furthermore position 46, 48 and 50 were preferred because, given previous experiments and the overall scores, the EGF C-terminus tail seems to play a critical role in the ligand function; see for example (57).

### Synthetic Peptides

The wild-type, N32R, D46T, K48T, and W50Y variants of EGF were ordered from Scrum Net Co with purity >90% and quantity 5 mg/mL. These peptides were used for ITC measurements, Circular Dichroism (CD) measurements, proliferation studies, and Western Blot (WB) analyses.

### Cell Lines

The *Bj5-tα human normal fibroblast cell line* was purchased from ATCC. Cells were grown in DMEM with 10% fetal bovine serum (FBS), and 5 μg/mL hygromycin B. The *Swiss Albino 3T3* mouse *normal fibroblast cell line* was obtained from the RIKEN Cell Bank. Cells were grown in DMEM, 10% FBS, 50 ug/mL gentamycin at 37°C in a 5% CO_2_ atmosphere with 95% humidity. The *A431 human epithelial carcinoma adherent cell line* (RIKEN Cell Bank) is a model skin cancer cell line with overexpressed EGFR used for oncogenic pathway studies (58).Cells were cultured in DMEM supplemented with 10% FBS (Sigma-Aldrich), 50 ug/mL gentamycin antibiotic or a combination of 100 unit/ml Penicillin G (Nacalai Tesque) and 100 μg/ml streptomycin sulfate (Nacalai Tesque). Experiments were conducted at 37°C in a 5% CO_2_-enriched air atmosphere with 95% humidity. Cell lines were grown and used for Western Blot and cell proliferation studies.

### Cell Proliferation Assay

We measured cell proliferation using a label-free, non-invasive, cellular confluence assay with IncuCyte Live-Cell Imaging Systems (Essen Bioscience). Human Bj5-tα (2,500 cells / well) and Mouse Swiss Albino 3T3 (1,000 cells/well) were seeded overnight on a 96-well plate (Corning) at 37°C in an incubator. The next day, cells were treated with WT EGF and mutants at 1 nM, 10 nM and 100 nM concentrations and placed in an XL-3 incubation chamber maintained at 37°C. The plate was scanned using a 4x objective at 2-hr intervals over 3 days. Cell confluence was measured using IncuCyte Analysis Software. The IncuCyte Analyzer gives real-time confluence data based on segmentation of high-definition phase-contrast images. Cell proliferation is shown as an increase in confluence rate relative to control.

### Apoptosis Assay

Experiments were performed with the A431 human cancer cell line. 5,000 cells/well were seeded on a 96-well plate (Corning) and incubated at 37°C for 24 hr. Media were replaced with fresh DMEM containing WT EGF, or EGF mutants at 1, 10, and 100 nM concentrations and fluorescent annexin V green reagent. Plates were pre-warmed prior to data acquisition to avoid condensation and expansion of the plate, which affect autofocus. Images were captured every 2 hrs (4x) for 3 days in the IncuCyte system. Cell proliferation is reported as in the previous assay.

### Statistics

Proliferation and apoptosis experiments were performed in duplicates. All results are shown as the mean±s.d. Raw data was analyzed by two-way ANOVA with 95% confidence level. The multiple test was corrected using Bonferroni post hoc test. Prism 8 software was used for statistical analysis. Asterisks in the pictures show P-values using GraphPad convention: 0.1234 > (ns), 0.0332 > (*), 0.0021 > (**), 0.0002 > (***), 0.0001 > (****).

### Isothermal Titration Calorimetry (ITC)

All ITC studies employed a MicroCal PEAQ-ITC System (Malvern). For titration, both EGFR ECD (Sigma-Aldrich) and EGF variants were dialyzed into the same reaction buffer Milli-Q H2O (22 μm) at 25°C. Each titration involved serial injections of 13 × 3 μL aliquots of EGF variants (200 μM) into a solution of EGFR ECD (20 μM) in the sample cell. In each case, the reference cell was filled with the same reaction buffer as the control to determine the heat upon binding of the two components. The measured heat constant value was subtracted from the heat per injection prior to analysis of the data. The experiment was replicated twice. Results were analyzed by MicroCal PEAQ-ITC Analysis Software.

### Circular Dichroism (CD)

Far UV measurements were taken at a protein concentration of 0.1 μM, using a cuvette with a path length of 0.1 cm. Secondary structure content was calculated from far UV spectra using CAPITO software (59). Five scans in the 190-240-nm wavelength range were taken.

### Western Blot Analysis

A431 epidermoid carcinoma cells were harvested using Lysis Buffer (0.4% SDS, 1 mM DTT, 1%). Samples were incubated at 37°C for 10 min with Benzonase and centrifuged at 15,000 rpm at 22°C for 10 min. Supernatants were used for further analysis. Sample concentrations were measured with a BCA protein assay kit (ThermoFisher Scientific). Lysate were mixed with 4x Sample Loading Laemmli Buffer and incubated at 90°C for 5 min. Equal amounts of protein were loaded in 12% Mini PROTEAN^®^ TGX™ SDS-PAGE gel (Bio-Rad) and transferred to PVDF membranes (Trans-Blot Turbo RTA Mini 0.2 μm PVDF Transfer Kit). Membranes were blocked for 10 min with Bullet Blocking One (Nacalai) and reacted with monoclonal rabbit anti-EGFR antibody (Cell Signaling Technology, Inc.), Phospho-EGF Receptor (Tyr1173) (Cell Signaling Technology, Inc.), and rabbit anti-α-tubulin pAb (MBL) at dilution of 1:1000. Samples were incubated with Goat Anti-Rabbit IgG HRP at a 1:5000 dilution and chemiluminescent signals were detected by CDP Plus (ThermoFisher Scientific) and ChemiDoc touch MP (Bio-rad).

### Cross-linking assay

A431 cells were cultured in 6 well-plate to sub-confluency. The cells were starved for 16 h. After activation with EGF and EGF mutants for 30 min at 4°C, the cells were washed 3 times by ice-cold PBS. The cross-linking reaction was performed as previously reported (60). Briefly, crosslinking reagents bis(sulfosuccinimidyl)suberate (BS3) (Dojindo) were then added to a final concentration of 3.0 mM in PBS, and the reaction was incubated on ice for 15min. The reaction was quenched by further incubation with 250mM Glycine in PBS and incubated for 15 min on ice. The cells were washed 3 times by ice-cold PBS, and then lysed with 1% SDS in PBS containing proteinase inhibitor cocktail (Nacalai) on ice. The EGFR dimerization was analyzed by SDS-PAGE and western blotting.

### Microscale Thermophoresis

Recombinant human EGFR Protein (ECD, His Tag) was purchased from Sino Biological Inc. (Cosmo bio, Japan). The protein was labeled with Large Volume His-Tag Labeling Kit RED-tris-NTA 2nd Generation (Nanotemper, Munich, Germany) and diluted to 200 nM with 0.05% tween-PBS. EGF WT, N32R, D46T, K48T, W50Y, BSA were prepared by 2-fold serial dilution with 0.05% tween-PBS (4000 nM-0.122 nM). The EGFR and ligands were mixed 1:1 and incubated at room temperature for 5 min, and then loaded into standard capillaries. Microscale thermophoresis measurements were performed by using Monolith NT.115 (Nanotemper, Munich, Germany).

### Molecular Dynamics

All the simulations were performed with Gromacs version 2020 (61) using charmm36-mar2019 force field (62) and SPC216 for water. PDB ID 5wb7 was used as model and template. EGF structure model was extracted from PDB ID 1ivo and superimposed to the ligand using UCSF Chimera Match-Maker algorithm (63). Mutants were generated using Swissmodel webserver (64), while missing atoms were compensated using Scwrl4 (65). A system was composed of EGFR dimer in complex with one or two EGF wild type (WT), N32R, D46T, K48T or W50Y were solvated and neutralized using NaCl ions in a dodecahedral box. Then, energy temperature and pressure equilibrations were performed to the system following the guidelines in Lemkul 2019 (66).

A 100 ns production simulation was run using the Verlet cut-off scheme (67) for non-bonded interactions and LINCS as constraint algorithm (68). All the simulations reached convergence of RMSD. Long range electrostatic interactions were computed using the Particle Mesh Ewald method (69) using a dedicated GPU. We checked for differences in relative motions between the three simulations by extracting and concatenating the backbone trajectories using catDCD plugin of VMD (70), then performing a PCA using Bio3d R package (71).

## Data availability

The source code and alignments data to reproduce the analysis of this paper are available at https://zenodo.org/badge/latestdoi/287415954.

## Acknowledgments

We thank Tadashi Yamamoto, Ichiro Maruyama for providing the antibodies, Keiko Kono for the technical help with the WB experiments. We are grateful for the help and support provided by the Scientific Computing section of Research Support Division and in particular Jan Moren at OIST. We thank Madhuri Gade, Mirco Dindo, Harry Wilson and Benjamin Clifton for critical reading. We thank Pierre Matricon for the discussion on the molecular dynamics data, Russel Alenton on the cell proliferation assay and Christine Guzman on the western blot analyses, and Saacnicteh Toledo Patino for the help with the NanoTemper instrument.

## Author contributions

PL, SP, DM conceived and designed the project; SP developed and performed the computational models and analyses; DM and GU performed the experiments; all authors analyzed the data and wrote the manuscript.

## Funding and additional information

Financial support by the Okinawa Institute of Science and Technology to P.L. is gratefully acknowledged.

## Abbreviations

EGF: Epidermal Growth Factor;
EGFR: Epidermal Growth Factor Receptor;
HBEGF: Heparin-Binding Epidermal Growth Factor;
EPGN: Epigen;
BTC: Betacellulin;
EREG: Epiregulin;
AREG: Amphiregulin;
PPI: Protein-Protein Interaction;
MSA: Multiple Sequence Alignment;
MSTA: Multiple STructural Alignment;
ITC: Isothermal Titration Calorimetry;
MST: MicroScale thermophoresis;
DMEM: Dulbecco’s Modified Eagle Medium;
ECD: Extra Cellular Domain;
WT: Wild Type;
CD: circular dichroism

