## Supplementary Data for "Binding of single-mutant epidermal growth factor (EGF) ligands alter the stability of the EGF receptor dimer and promote growth signaling"

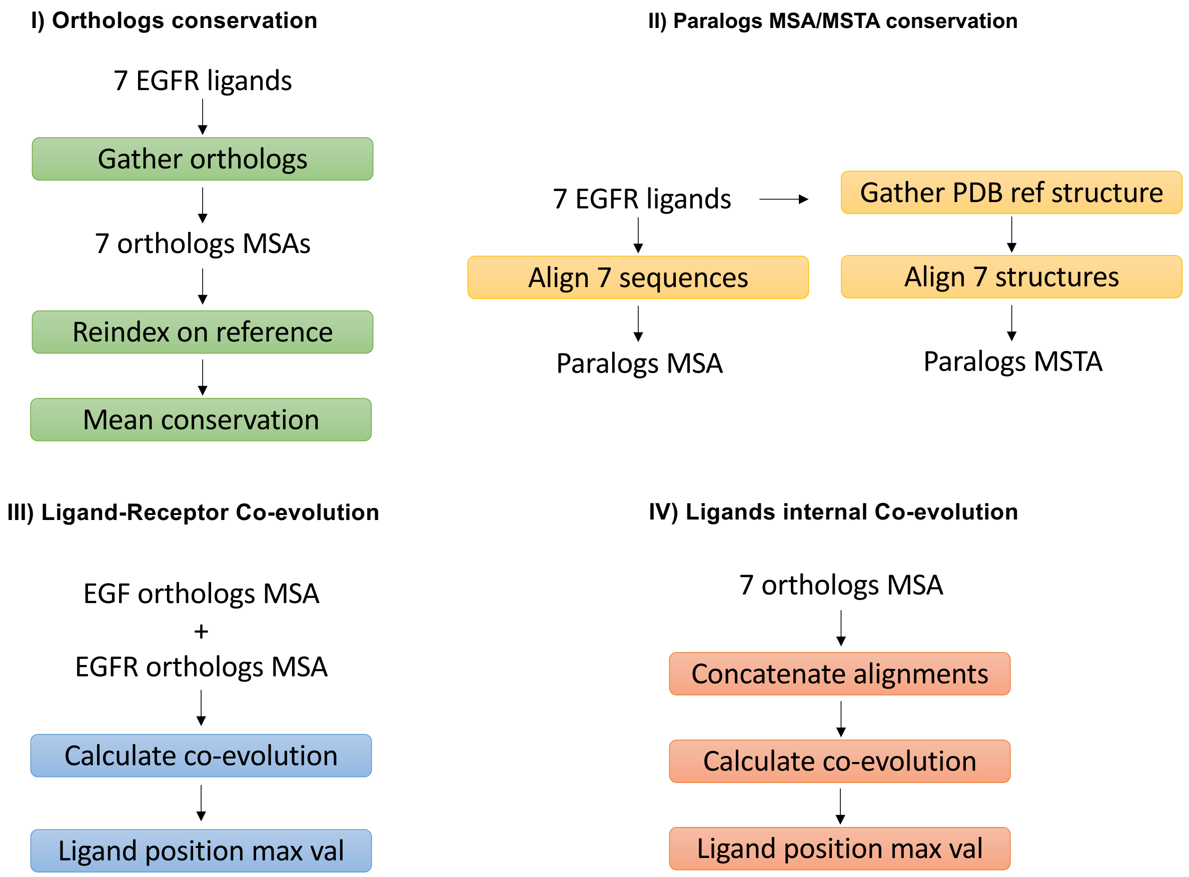

**Figure S1. DIRpred schema.** A schematic representation of DIRPred, as shown in Figure 7.

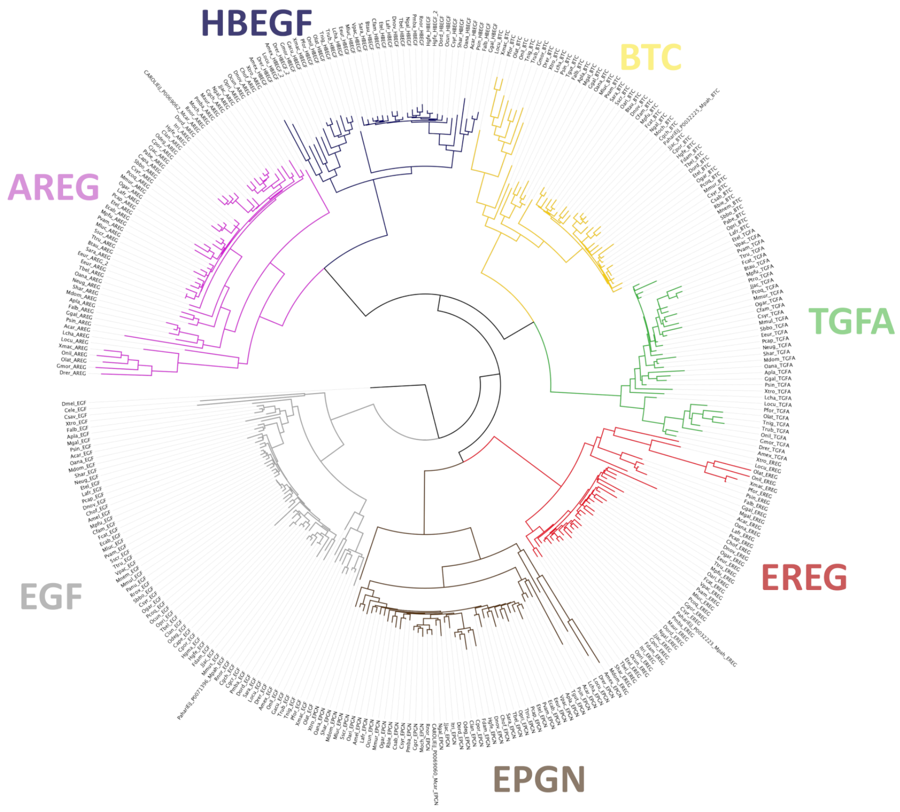

**Figure S2. EGFR ligands orthologs tree.** The tree was obtained using Maximum Likelihood algorithm. *C elegans* EGF was used as outgroup. The EGFR ligands cluster neatly. Interestingly, fish ligands are found to have longer branches on average. The tree was made using iqtree ModelFinder. The best-fit model used is JTT+R6.

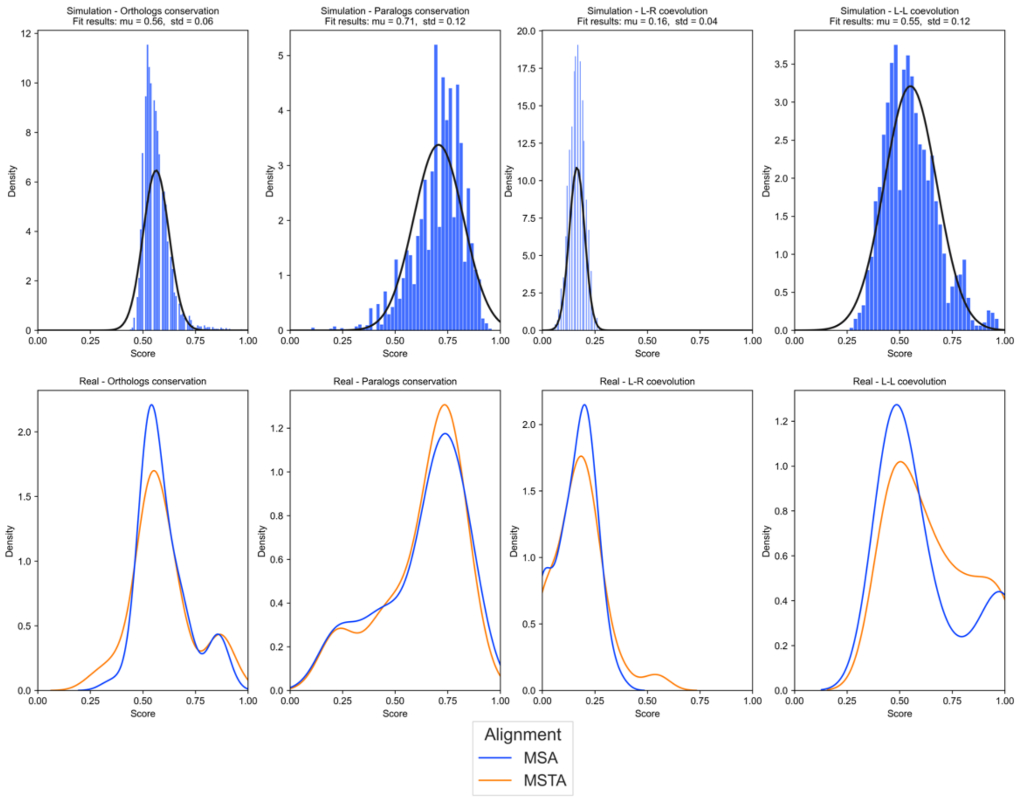

**Figure S3. DIRpred statistics.** The four columns represent the four DIRpred scores. The top row shows the score distributions of the DIRpred analysis of a dataset of 100 repetitions of simulated evolution. The simulations started from a random 53 amino acid long sequence and were carried out using the WAG model. On the bottom row, the distribution of scores from the real alignments of EGF and its homologs, divided by the sequence or structural alignment seed used for the analysis.

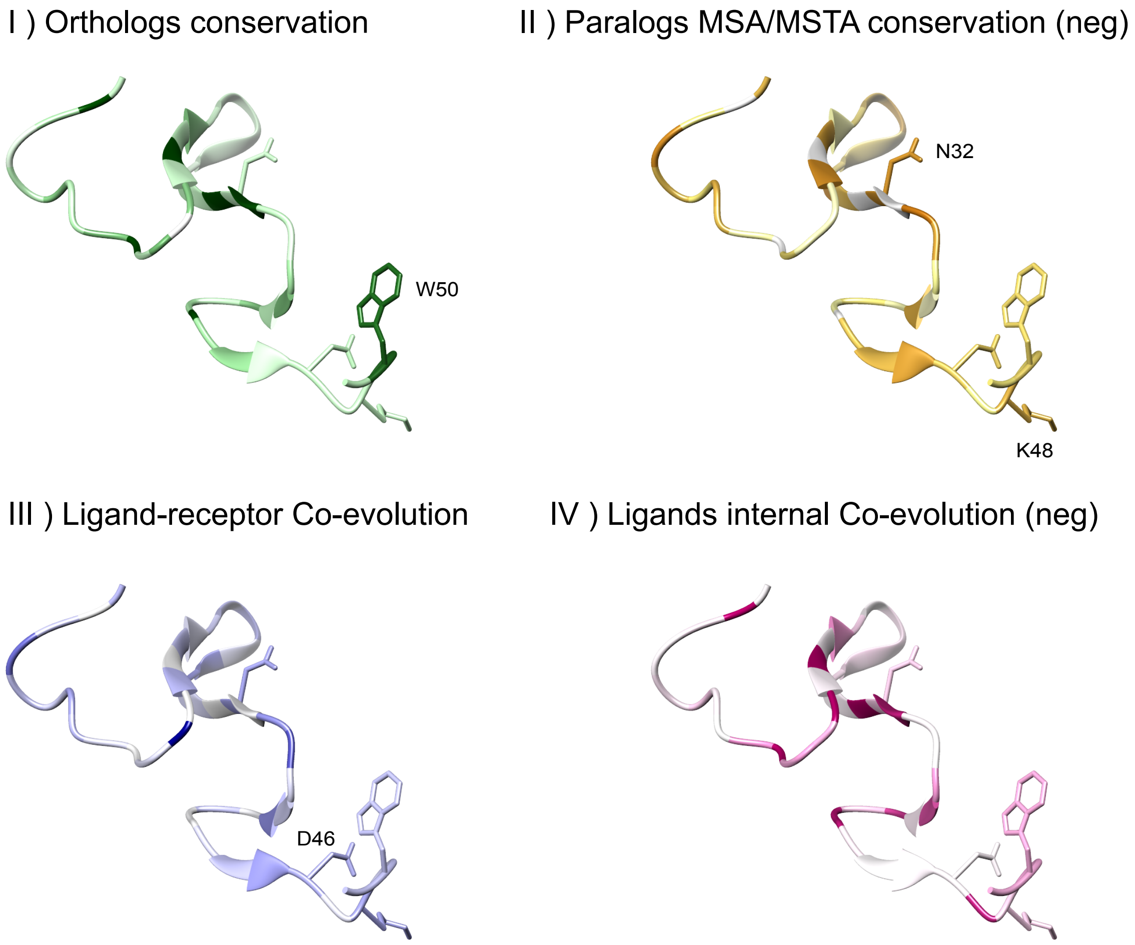

**Figure S4. DIRpred score.** Structure of EGF (from PDB: 1IVO) colored by the four individual scores. The four sites chosen for mutation are shown in stick representation.

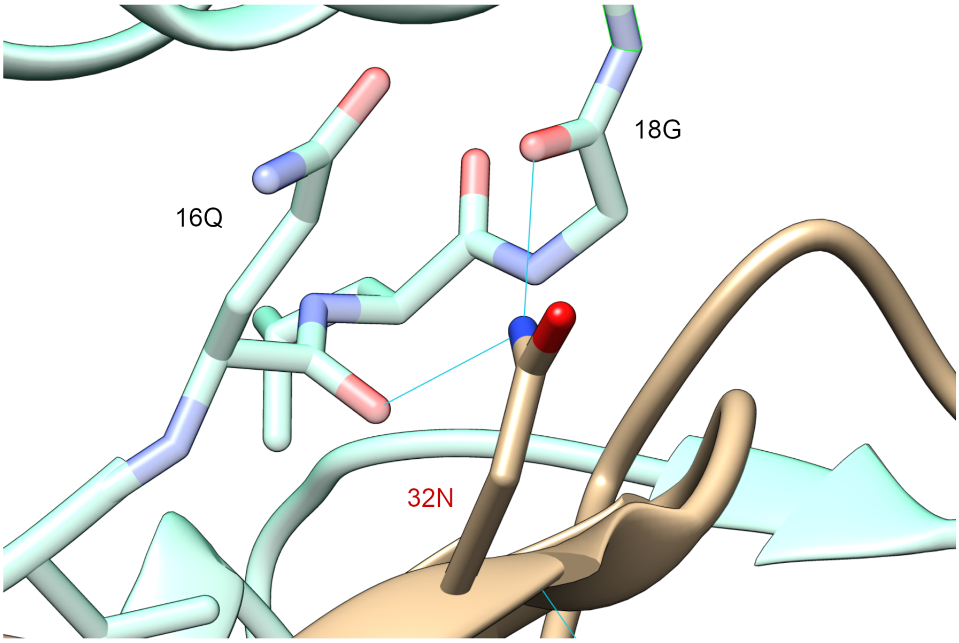

**Figure S5. Crystalized structure of WT EGF with the highlighted position N32, In the interaction with the position Q16 and G18 of the EGFR receptor backbone.**

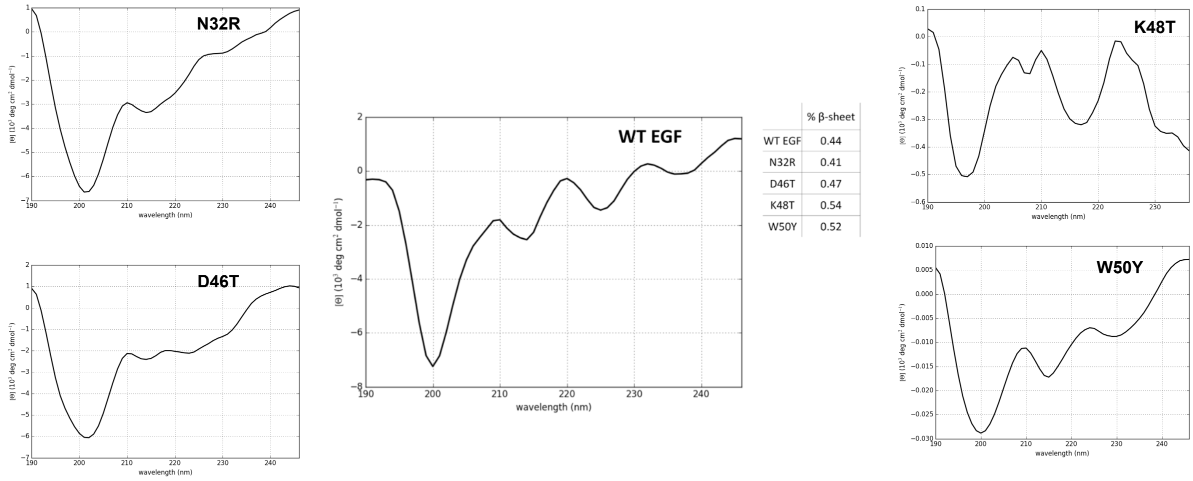

| | % β-sheet |
|---|---|
| WT EGF | 0.44 |
| N32R | 0.41 |
| D46T | 0.47 |
| K48T | 0.54 |
| W50Y | 0.52 |

**Figure S6. The validation of ordered structure of WT EGF and variants.** 0.2 mg/mL of the sample was added to a 10 µm cuvette and analyzed by circular dichroism spectroscopy. All samples showed the presence of a comparable amount of $\beta$-sheet. The CD spectrum of EGF was similar to previously published results (26). The plot and analysis was made using CAPITO web tool (27).

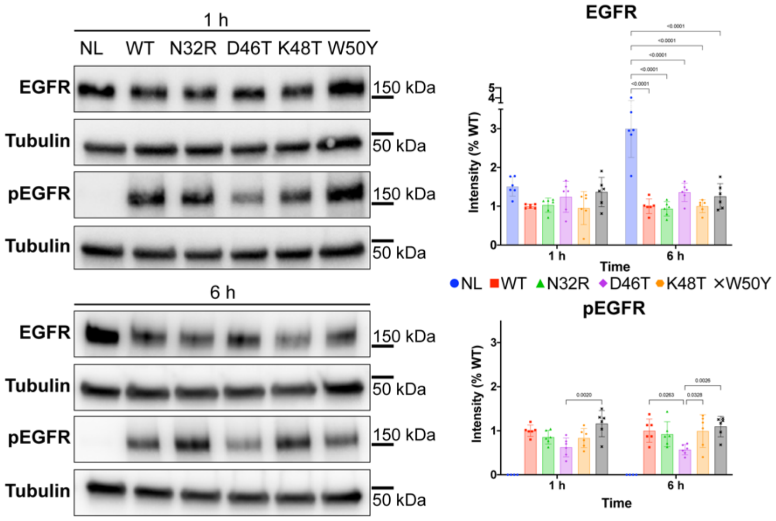

**Figure S7. Effects of WT and single-mutant EGF on the receptor phosphorylation and dimerization in A431 cells at additional time points.** The effects on the amount of EGFR and the phosphorylation level of Tyr-1173 after treating A431 cells with 100nM concentration of different ligands at 1 and 6 h. The membrane containing EGFR and tubulin was separated after the transfer. The WB was performed in the same conditions as in Figure 4. Data is shown of at least four biological duplicates. The bars represent mean ± s.d. of at least four biological repeats. The number on top of the bars shows the p-values of a 2-way ANOVA multiple comparison corrected for multiple sampling using the Bonferroni correction. Band intensity estimates were calculated using BIO-RAD ImageLab software (BioRad). Plots and statistics were performed using PRISM software (GraphPad). Details of the ANOVA are found in Table S3.

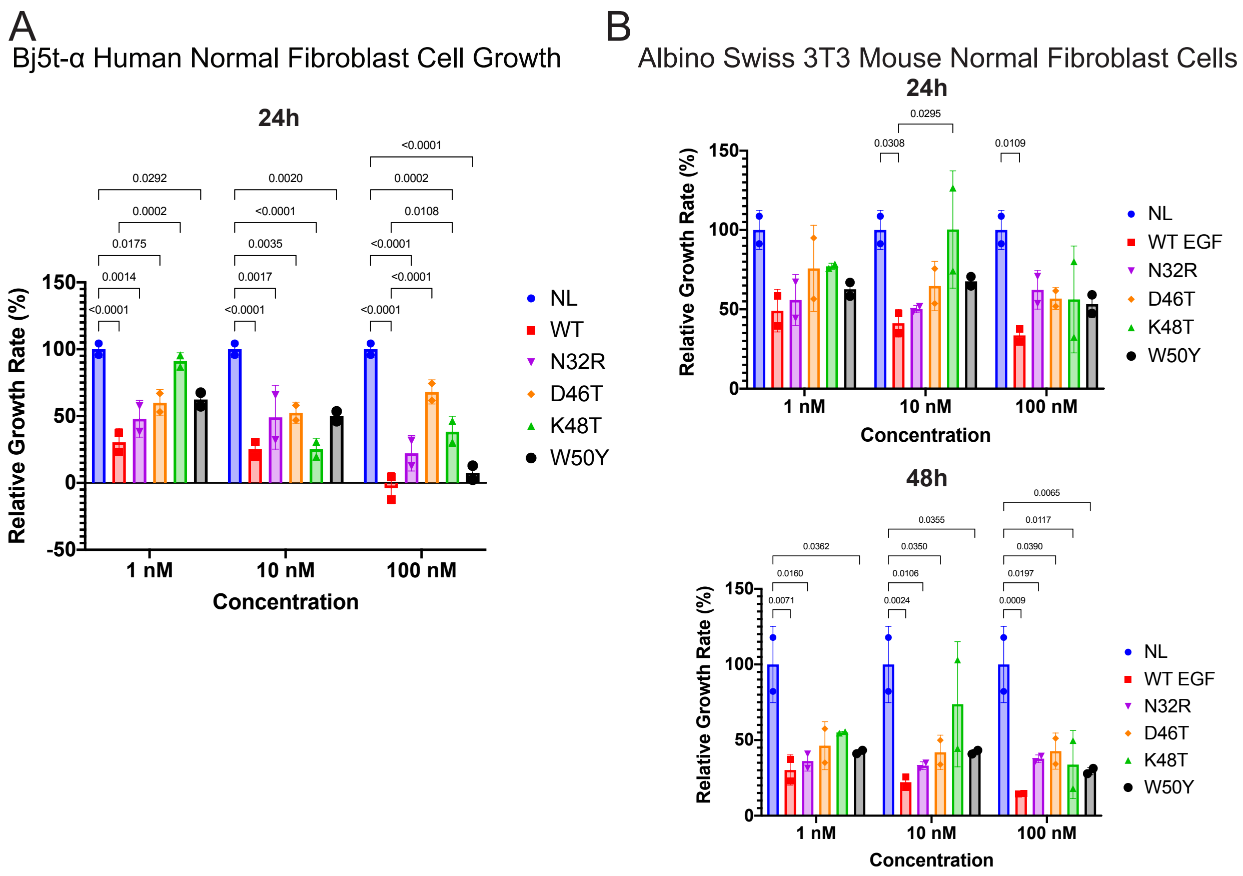

**Figure S8. Effect of 1nM, 10nM and 100nM concentrations of EGF variants on the proliferation of the human normal fibroblast Bj5-tα cell line and mouse normal fibroblast Albino Swiss 3T3**

**cell line.** The data represent growth as a percentage of the control based on the confluence of cells (mean ± standard deviation) for each concentration of EGF. The percentage confluence was estimated 24 h and 48 h after the treatment (two replicates per treatment). Details of the ANOVA are found in Table S3.

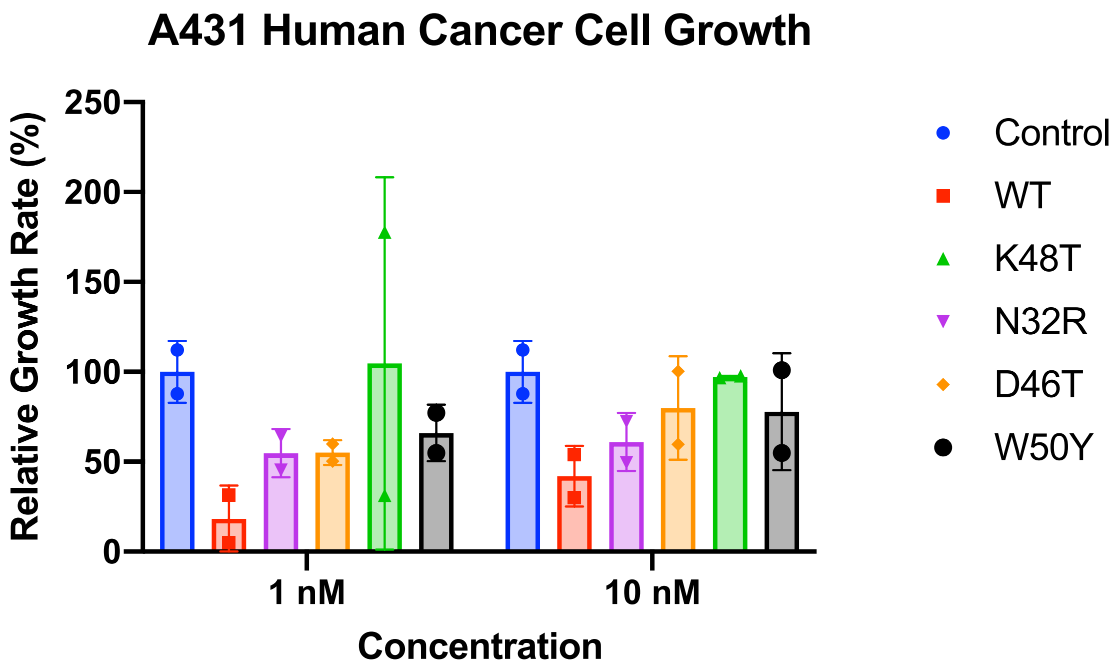

**Figure S9. Effect of 1nM and 10nM concentration of EGF variants on the proliferation of A431 skin cancer cells.** The data represent the relative growth percentage to control based on the confluence of cells (mean ± standard deviation) for each concentration of EGF variants. The percentage confluence was estimated 48 h after the treatment (two replicates/treatment).

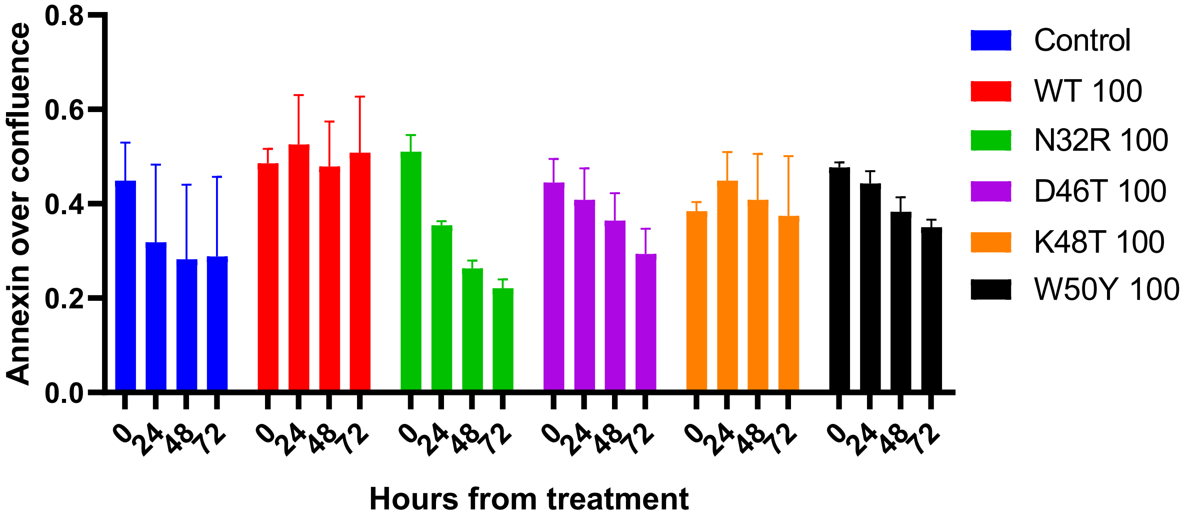

**Figure S10. A431 cell apoptotic assay.** After 24h growth incubation, A431 cells were treated with saturating concentrations of WT or mutant EGF. The green reagent Annexin V confluence was related to the phase confluence (cell growth). WT EGF is the only sample to show a stable trend after several days, while all other samples show a reduction in the ratio of Annexin signal to confluence. The apparent high level of K48T could be explained by the change in morphology of the cells.

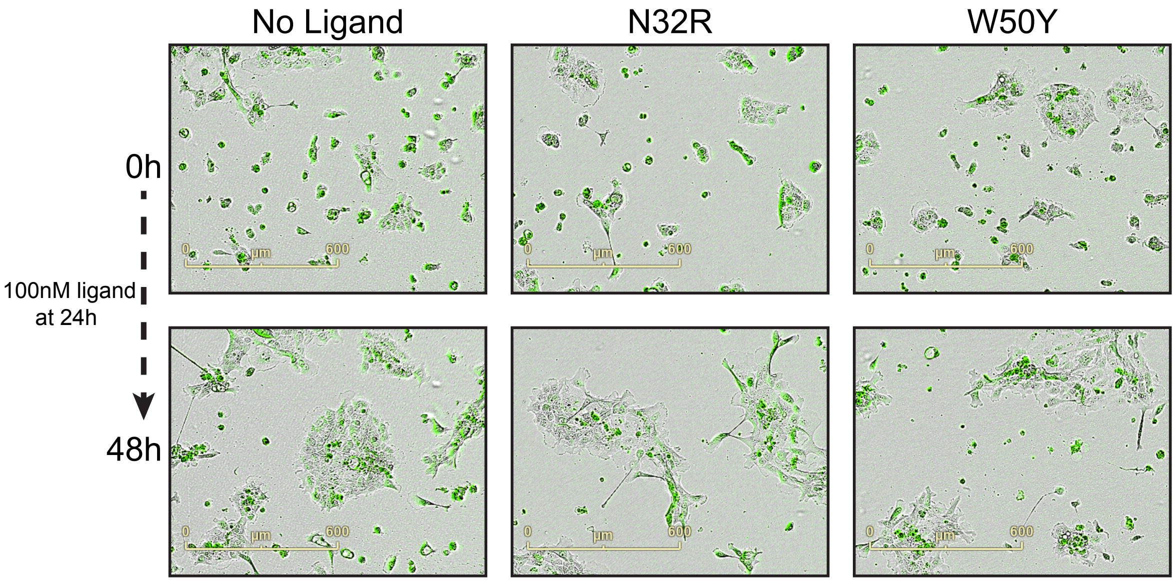

**Figure S11. Growth assay of A431 cells.** Comparison of A431 cell growth after treatment with 100 nM EGF variants N32R and W50Y or untreated. Cells were labeled with fluorescent Annexin V Green Reagent. Plates were pre-warmed prior to data acquisition to avoid condensation and expansion of the plate, which affect autofocus. Images were captured every 2 h (4x magnification) for 3 days in the IncuCyte system.

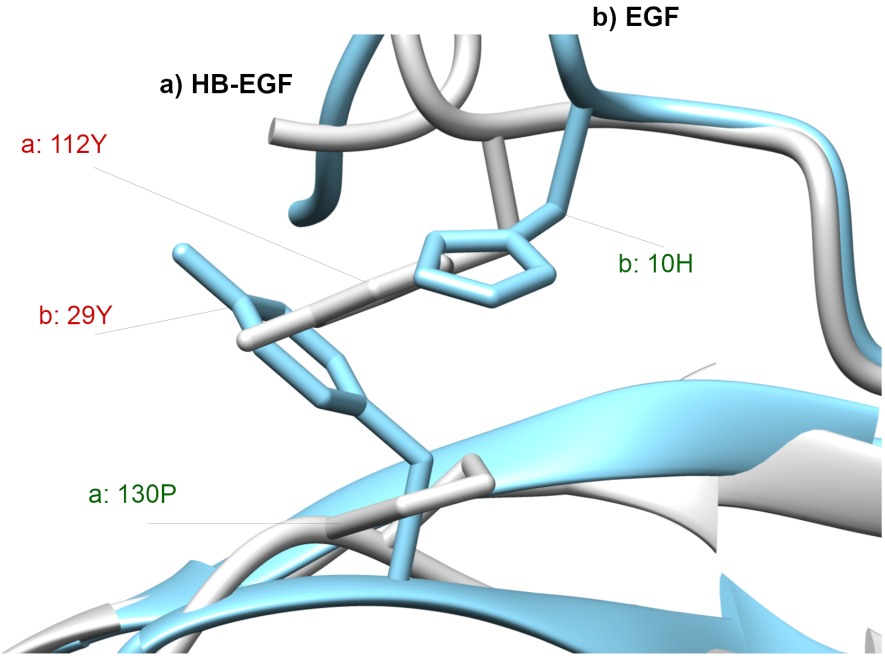

**Figure S12. Highlight on the EGF-specific residue swapping problem.** EGF is colored in teal, while HB-EGF is colored in white. The tyrosine in EGF position 29 (29Y) is not found on the corresponding position of HB-EGF. Instead, a proline is found (130P). Though, the interaction between the two residue is probably maintained because, in place of the histidine 10 (10H) of EGF, HB-EGF shows a tyrosine (112Y). When this type of residue swapping is ligand-specific, the lowering of the conservation measures over-estimates the real change happening in that position.

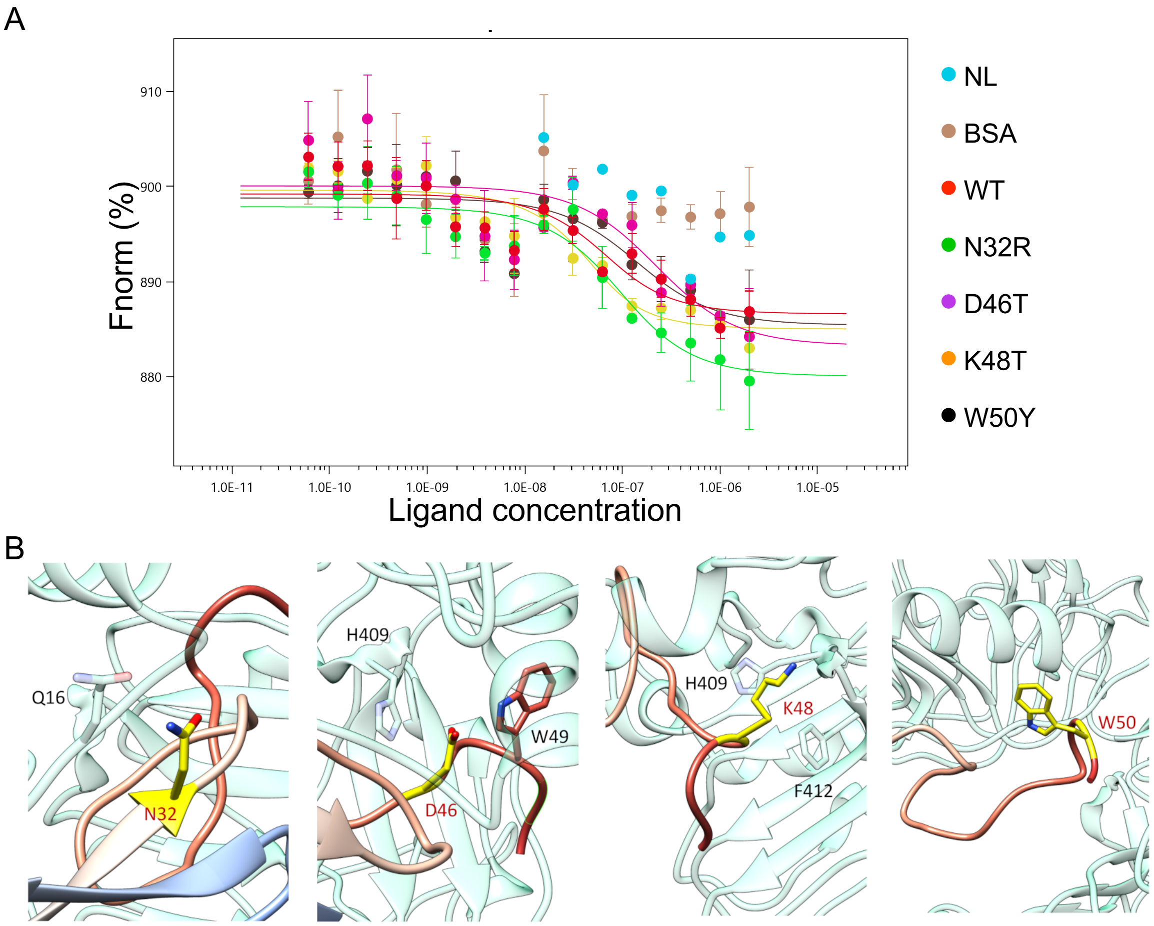

**Figure S13. Details of the mutants and their binding.** (A) The MTS experiment including the No Ligand (NL) sample. BSA shows a response within the NL range, while all other ligands show a transition at a similar concentration towards higher fluorescent signal. (B) Zoom-in of the four mutations included in a model starting from PDBID:1IVO.

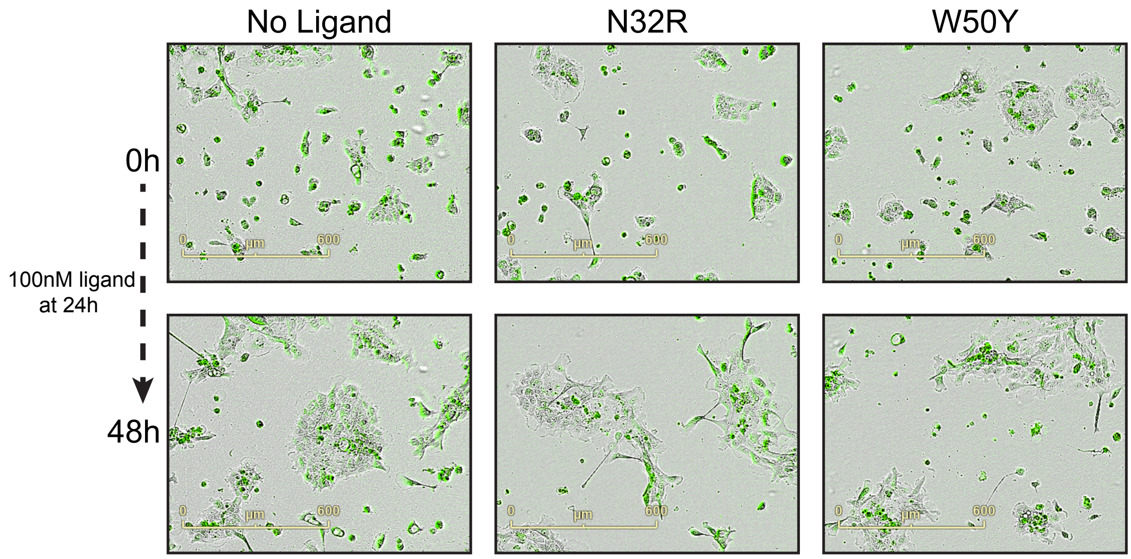

**Figure S14. A431 cells apoptosis experiment comparison for No Ligand, N32R, and W50Y samples.** Comparison of A431 cell growth after treatment with NL (No ligand), 100 nM N32R or W50Y samples. Cells were labeled with fluorescent Annexin V Green Reagent. Plates were pre-warmed prior to data acquisition to avoid condensation and expansion of the plate, which affect autofocus. Images were captured every 2 h (4x magnification) for 3 days in the IncuCyte system.

**Table S1. DIRpred score of each position of EGF.** The table resembles the actual output of the DIRpred pipeline, reporting the four individual scores and the combined score. Furthermore, we added the P-value associated to each individual score and the sum of P-value ranking.

| | POS | ORTHOLOGS CONSERVATION MSA | P-VALUE ORTHOLOGS CONSERVATION MSA | PARALOGS CONSERVATION MSA | P-VALUE PARALOGS CONSERVATION MSA | L-R COEVOLUTION MSA | P-VALUE L-R COEVOLUTION MSA | L-L COEVOLUTION MSA | P-VALUE L-L COEVOLUTION MSA | ORTHOLOGS CONSERVATION MSTA | P-VALUE ORTHOLOGS CONSERVATION MSTA | PARALOGS CONSERVATION MSTA | P-VALUE PARALOGS CONSERVATION MSTA | L-R COEVOLUTION MSTA | P-VALUE L-R COEVOLUTION MSTA | L-L COEVOLUTION MSTA | P-VALUE L-L COEVOLUTION MSTA | AVG DIR SCORE | P-VALUE SUM | P-VALUE RANK | MSA P-VALUE SUM |
|---|---|---|---|---|---|---|---|---|---|---|---|---|---|---|---|---|---|---|---|---|---|
| 1 | 1N | 0.55 | 0.58 | 0.25 | 0.36 | 0.18 | 0.34 | 0.56 | 0.81 | 0.59 | 0.33 | 0.27 | 0.42 | 0.24 | 0.02 | 0.24 | 0.05 | 0.53 | 2.90 | 5 | 2.09 |
| 2 | 2S | 0.59 | 0.33 | 0.27 | 0.42 | 0.20 | 0.17 | 0.59 | 0.87 | 0.41 | 0.99 | 0.27 | 0.42 | 0.39 | 0.00 | 0.36 | 0.24 | 0.51 | 3.44 | 12 | 1.79 |
| 3 | 3D | 0.53 | 0.70 | 0.15 | 0.11 | 0.20 | 0.17 | 0.59 | 0.87 | 0.26 | 1.00 | 0.27 | 0.42 | 0.57 | 0.00 | 0.47 | 0.57 | 0.51 | 3.84 | 24 | 1.85 |
| 4 | 4S | 0.55 | 0.58 | 0.32 | 0.59 | 0.25 | 0.01 | 0.52 | 0.72 | 0.31 | 1.00 | 0.27 | 0.42 | 0.51 | 0.00 | 0.47 | 0.57 | 0.50 | 3.88 | 25 | 1.90 |
| 5 | 5E | 0.51 | 0.80 | 0.25 | 0.36 | 0.21 | 0.11 | 0.52 | 0.72 | 0.53 | 0.70 | 0.28 | 0.45 | 0.18 | 0.34 | 0.58 | 0.85 | 0.48 | 4.33 | 45 | 1.98 |
| 6 | 6C | 0.86 | 0.00 | 0.78 | 1.00 | 0.00 | 1.00 | 0.00 | 0.00 | 0.86 | 0.00 | 0.77 | 1.00 | 0.00 | 1.00 | 0.00 | 0.00 | 0.52 | 4.00 | 29 | 2.00 |
| 7 | 7P | 0.57 | 0.45 | 0.31 | 0.55 | 0.16 | 0.55 | 0.53 | 0.74 | 0.61 | 0.22 | 0.40 | 0.82 | 0.18 | 0.34 | 0.46 | 0.54 | 0.48 | 4.21 | 43 | 2.30 |
| 8 | 8L | 0.48 | 0.91 | 0.13 | 0.08 | 0.25 | 0.01 | 0.53 | 0.74 | 0.40 | 1.00 | 0.18 | 0.17 | 0.29 | 0.00 | 0.54 | 0.77 | 0.51 | 3.68 | 18 | 1.75 |
| 9 | 9S | 0.56 | 0.52 | 0.30 | 0.52 | 0.18 | 0.34 | 0.47 | 0.57 | 0.56 | 0.52 | 0.27 | 0.42 | 0.18 | 0.34 | 0.53 | 0.74 | 0.49 | 3.96 | 27 | 1.94 |
| 10 | 10H | 0.68 | 0.03 | 0.43 | 0.88 | 0.12 | 0.89 | 0.40 | 0.35 | 0.64 | 0.10 | 0.25 | 0.36 | 0.15 | 0.66 | 0.46 | 0.54 | 0.51 | 3.79 | 23 | 2.14 |
| 11 | 11D | 0.53 | 0.70 | 0.26 | 0.39 | 0.25 | 0.01 | 0.47 | 0.57 | 0.57 | 0.45 | 0.23 | 0.29 | 0.18 | 0.34 | 0.58 | 0.85 | 0.50 | 3.60 | 15 | 1.67 |
| 12 | 12G | 0.60 | 0.27 | 0.28 | 0.45 | 0.19 | 0.24 | 0.56 | 0.81 | 0.57 | 0.45 | 0.28 | 0.45 | 0.21 | 0.11 | 0.58 | 0.85 | 0.48 | 3.65 | 17 | 1.78 |
| 13 | 13Y | 0.68 | 0.03 | 0.55 | 0.98 | 0.09 | 0.98 | 0.19 | 0.02 | 0.68 | 0.03 | 0.50 | 0.96 | 0.10 | 0.96 | 0.35 | 0.21 | 0.50 | 4.17 | 40 | 2.01 |
| 14 | 14C | 0.86 | 0.00 | 0.78 | 1.00 | 0.00 | 1.00 | 0.00 | 0.00 | 0.86 | 0.00 | 0.78 | 1.00 | 0.00 | 1.00 | 0.00 | 0.00 | 0.52 | 4.00 | 30 | 2.00 |
| 15 | 15L | 0.54 | 0.64 | 0.38 | 0.77 | 0.09 | 0.98 | 0.35 | 0.21 | 0.54 | 0.64 | 0.38 | 0.77 | 0.09 | 0.98 | 0.33 | 0.17 | 0.48 | 5.16 | 51 | 2.60 |
| 16 | 16H | 0.70 | 0.01 | 0.57 | 0.99 | 0.10 | 0.96 | 0.35 | 0.21 | 0.70 | 0.01 | 0.57 | 0.99 | 0.10 | 0.96 | 0.24 | 0.05 | 0.48 | 4.19 | 41 | 2.18 |
| 17 | 17D | 0.35 | 1.00 | 0.27 | 0.42 | 0.36 | 0.00 | 0.35 | 0.21 | 0.35 | 1.00 | 0.27 | 0.42 | 0.36 | 0.00 | 0.35 | 0.21 | 0.52 | 3.27 | 10 | 1.63 |
| 18 | 18G | 0.66 | 0.06 | 0.61 | 1.00 | 0.00 | 1.00 | 0.00 | 0.00 | 0.66 | 0.06 | 0.61 | 1.00 | 0.00 | 1.00 | 0.00 | 0.00 | 0.51 | 4.11 | 37 | 2.05 |
| 19 | 19V | 0.52 | 0.76 | 0.18 | 0.17 | 0.20 | 0.17 | 0.53 | 0.74 | 0.52 | 0.76 | 0.18 | 0.17 | 0.20 | 0.17 | 0.54 | 0.77 | 0.50 | 3.69 | 19 | 1.83 |
| 20 | 20C | 0.86 | 0.00 | 0.78 | 1.00 | 0.00 | 1.00 | 0.00 | 0.00 | 0.86 | 0.00 | 0.78 | 1.00 | 0.00 | 1.00 | 0.00 | 0.00 | 0.52 | 4.00 | 30 | 2.00 |
| 21 | 21M | 0.54 | 0.64 | 0.23 | 0.29 | 0.17 | 0.44 | 0.58 | 0.85 | 0.54 | 0.64 | 0.23 | 0.29 | 0.17 | 0.44 | 0.51 | 0.69 | 0.48 | 4.30 | 44 | 2.23 |
| 22 | 22Y | 0.68 | 0.03 | 0.55 | 0.98 | 0.09 | 0.98 | 0.28 | 0.09 | 0.68 | 0.03 | 0.55 | 0.98 | 0.09 | 0.98 | 0.22 | 0.03 | 0.49 | 4.11 | 36 | 2.08 |
| 23 | 23I | 0.53 | 0.70 | 0.37 | 0.74 | 0.19 | 0.24 | 0.46 | 0.54 | 0.53 | 0.70 | 0.37 | 0.74 | 0.19 | 0.24 | 0.36 | 0.24 | 0.49 | 4.15 | 38 | 2.22 |
| 24 | 24E | 0.53 | 0.70 | 0.27 | 0.42 | 0.23 | 0.04 | 0.57 | 0.84 | 0.55 | 0.58 | 0.33 | 0.62 | 0.22 | 0.07 | 0.52 | 0.72 | 0.48 | 3.98 | 28 | 1.99 |
| 25 | 25A | 0.52 | 0.76 | 0.24 | 0.32 | 0.27 | 0.00 | 0.46 | 0.54 | 0.53 | 0.70 | 0.19 | 0.19 | 0.25 | 0.01 | 0.45 | 0.50 | 0.53 | 3.02 | 9 | 1.62 |
| 26 | 26L | 0.52 | 0.76 | 0.34 | 0.65 | 0.15 | 0.66 | 0.42 | 0.41 | 0.52 | 0.76 | 0.39 | 0.79 | 0.14 | 0.75 | 0.37 | 0.26 | 0.48 | 5.03 | 50 | 2.47 |
| 27 | 27D | 0.51 | 0.80 | 0.34 | 0.65 | 0.30 | 0.00 | 0.42 | 0.41 | 0.51 | 0.80 | 0.34 | 0.65 | 0.30 | 0.00 | 0.43 | 0.44 | 0.51 | 3.76 | 22 | 1.86 |
| 28 | 28K | 0.46 | 0.95 | 0.34 | 0.65 | 0.29 | 0.00 | 0.43 | 0.44 | 0.46 | 0.95 | 0.34 | 0.65 | 0.29 | 0.00 | 0.45 | 0.50 | 0.49 | 4.15 | 39 | 2.04 |
| 29 | 29Y | 0.62 | 0.18 | 0.13 | 0.08 | 0.15 | 0.66 | 0.53 | 0.74 | 0.62 | 0.18 | 0.13 | 0.08 | 0.15 | 0.66 | 0.54 | 0.77 | 0.53 | 3.34 | 11 | 1.66 |
| 30 | 30A | 0.54 | 0.64 | 0.30 | 0.52 | 0.12 | 0.89 | 0.53 | 0.74 | 0.54 | 0.64 | 0.30 | 0.52 | 0.12 | 0.89 | 0.53 | 0.74 | 0.46 | 5.59 | 53 | 2.80 |
| 31 | 31C | 0.86 | 0.00 | 0.78 | 1.00 | 0.00 | 1.00 | 0.00 | 0.00 | 0.86 | 0.00 | 0.78 | 1.00 | 0.00 | 1.00 | 0.00 | 0.00 | 0.52 | 4.00 | 30 | 2.00 |
| 32 | 32N | 0.51 | 0.80 | 0.18 | 0.17 | 0.22 | 0.07 | 0.43 | 0.44 | 0.51 | 0.80 | 0.18 | 0.17 | 0.22 | 0.07 | 0.45 | 0.50 | 0.53 | 3.02 | 8 | 1.48 |
| 33 | 33C | 0.85 | 0.00 | 0.78 | 1.00 | 0.01 | 1.00 | 0.03 | 0.00 | 0.85 | 0.00 | 0.78 | 1.00 | 0.01 | 1.00 | 0.07 | 0.00 | 0.51 | 4.00 | 34 | 2.00 |
| 34 | 34V | 0.55 | 0.58 | 0.13 | 0.08 | 0.24 | 0.02 | 0.53 | 0.74 | 0.55 | 0.58 | 0.13 | 0.08 | 0.24 | 0.02 | 0.55 | 0.79 | 0.53 | 2.90 | 4 | 1.43 |
| 35 | 35V | 0.50 | 0.84 | 0.23 | 0.29 | 0.28 | 0.00 | 0.56 | 0.81 | 0.50 | 0.84 | 0.22 | 0.27 | 0.30 | 0.00 | 0.57 | 0.84 | 0.50 | 3.90 | 26 | 1.96 |
| 36 | 36G | 0.61 | 0.22 | 0.53 | 0.98 | 0.09 | 0.98 | 0.20 | 0.02 | 0.62 | 0.18 | 0.45 | 0.91 | 0.10 | 0.96 | 0.38 | 0.29 | 0.48 | 4.53 | 48 | 2.20 |
| 37 | 37Y | 0.70 | 0.01 | 0.67 | 1.00 | 0.10 | 0.96 | 0.08 | 0.00 | 0.65 | 0.08 | 0.59 | 0.99 | 0.16 | 0.55 | 0.22 | 0.03 | 0.51 | 3.63 | 16 | 1.97 |
| 38 | 38I | 0.52 | 0.76 | 0.29 | 0.49 | 0.27 | 0.00 | 0.50 | 0.66 | 0.56 | 0.52 | 0.24 | 0.32 | 0.22 | 0.07 | 0.55 | 0.79 | 0.50 | 3.60 | 14 | 1.90 |
| 39 | 39G | 0.66 | 0.06 | 0.61 | 1.00 | 0.01 | 1.00 | 0.02 | 0.00 | 0.63 | 0.14 | 0.55 | 0.98 | 0.04 | 1.00 | 0.20 | 0.02 | 0.50 | 4.20 | 42 | 2.05 |
| 40 | 40E | 0.53 | 0.70 | 0.37 | 0.74 | 0.18 | 0.34 | 0.48 | 0.60 | 0.54 | 0.64 | 0.40 | 0.82 | 0.18 | 0.34 | 0.34 | 0.19 | 0.48 | 4.36 | 46 | 2.38 |
| 41 | 41R | 0.60 | 0.27 | 0.55 | 0.98 | 0.01 | 1.00 | 0.02 | 0.00 | 0.60 | 0.27 | 0.54 | 0.98 | 0.01 | 1.00 | 0.06 | 0.00 | 0.51 | 4.51 | 47 | 2.26 |
| 42 | 42C | 0.86 | 0.00 | 0.78 | 1.00 | 0.01 | 1.00 | 0.02 | 0.00 | 0.86 | 0.00 | 0.77 | 1.00 | 0.01 | 1.00 | 0.06 | 0.00 | 0.51 | 4.00 | 33 | 2.00 |
| 43 | 43Q | 0.61 | 0.22 | 0.32 | 0.59 | 0.12 | 0.89 | 0.45 | 0.50 | 0.61 | 0.22 | 0.31 | 0.55 | 0.12 | 0.89 | 0.57 | 0.84 | 0.48 | 4.70 | 49 | 2.20 |
| 44 | 44Y | 0.60 | 0.27 | 0.26 | 0.39 | 0.20 | 0.17 | 0.56 | 0.81 | 0.66 | 0.06 | 0.26 | 0.39 | 0.16 | 0.55 | 0.57 | 0.84 | 0.50 | 3.47 | 13 | 1.64 |
| 45 | 45R | 0.51 | 0.80 | 0.18 | 0.17 | 0.22 | 0.07 | 0.57 | 0.84 | 0.51 | 0.80 | 0.26 | 0.39 | 0.24 | 0.02 | 0.50 | 0.66 | 0.50 | 3.74 | 21 | 1.87 |
| 46 | 46D | 0.56 | 0.52 | 0.34 | 0.65 | 0.20 | 0.17 | 0.53 | 0.74 | 0.54 | 0.64 | 0.29 | 0.49 | 0.22 | 0.07 | 0.55 | 0.79 | 0.48 | 4.07 | 35 | 2.08 |
| 47 | 47L | 0.52 | 0.76 | 0.48 | 0.94 | 0.08 | 0.99 | 0.06 | 0.00 | 0.52 | 0.76 | 0.47 | 0.93 | 0.06 | 1.00 | 0.06 | 0.00 | 0.51 | 5.37 | 52 | 2.69 |
| 48 | 48K | 0.55 | 0.58 | 0.24 | 0.32 | 0.22 | 0.07 | 0.50 | 0.66 | 0.53 | 0.70 | 0.22 | 0.27 | 0.23 | 0.04 | 0.38 | 0.29 | 0.52 | 2.92 | 6 | 1.63 |
| 49 | 49W | 0.66 | 0.06 | 0.15 | 0.11 | 0.13 | 0.83 | 0.56 | 0.81 | 0.93 | 0.00 | 0.20 | 0.21 | 0.13 | 0.83 | 0.10 | 0.00 | 0.60 | 2.86 | 3 | 1.81 |
| 50 | 50W | 0.73 | 0.00 | 0.17 | 0.15 | 0.20 | 0.17 | 0.53 | 0.74 | 0.89 | 0.00 | 0.17 | 0.15 | 0.15 | 0.66 | 0.15 | 0.01 | 0.62 | 1.87 | 1 | 1.06 |
| 51 | 51E | 0.53 | 0.70 | 0.26 | 0.39 | 0.17 | 0.44 | 0.56 | 0.81 | 0.56 | 0.52 | 0.27 | 0.42 | 0.17 | 0.44 | 0.14 | 0.01 | 0.52 | 3.73 | 20 | 2.34 |

| | | | | | | | | | | | | | | | | | | | | |
|---|---|---|---|---|---|---|---|---|---|---|---|---|---|---|---|---|---|---|---|---|
| **52** | 52L | 0.54 | 0.64 | 0.20 | 0.21 | 0.23 | 0.04 | 0.53 | 0.74 | 0.48 | 0.91 | 0.27 | 0.42 | 0.25 | 0.01 | 0.19 | 0.02 | 0.54 | 3.00 | 7 | 1.64 |
| **53** | 53R | 0.55 | 0.58 | 0.15 | 0.11 | 0.21 | 0.11 | 0.57 | 0.84 | 0.53 | 0.70 | 0.27 | 0.42 | 0.25 | 0.01 | 0.26 | 0.06 | 0.54 | 2.83 | 2 | 1.64 |

**Table S2. EGF mutants relative ranking for the DIRpred score.** The four partial scores were ranked from 1st to 55th on each protein site, for the two methods using either the Multiple Sequence Alignment (MSA) or the Multiple Structure Alignment (MSTA). The ranking of the four sites chosen for mutagenesis are reported.

**Asn-32:**

| *Score* | *I* | *II* | *III* | *IV* |
|---|---|---|---|---|
| *MSA* | 46 | 8 | 13 | 21 |
| *MSTA* | 44 | 5 | 17 | 31 |

**Asp-46:**

| *Score* | *I* | *II* | *III* | *IV* |
|---|---|---|---|---|
| *MSA* | 24 | 31 | 18 | 33 |
| *MSTA* | 32 | 28 | 17 | 46 |

**Lys-48:**

| *Score* | *I* | *II* | *III* | *IV* |
|---|---|---|---|---|
| *MSA* | 26 | 14 | 13 | 29 |
| *MSTA* | 37 | 9 | 15 | 27 |

**Trp-50:**

| *Score* | *I* | *II* | *III* | *IV* |
|---|---|---|---|---|
| *MSA* | 7 | 7 | 18 | 33 |
| *MSTA* | 2 | 3 | 33 | 12 |

**Table S3. ANOVA details.** The full results of the ANOVA analyses. When not specified, the row factor represent the times of dosage for the treatment, while the column factor represent the ligands used in the treatment.

| Figure 4A | | | | | |
|---|---|---|---|---|---|
| **Source of Variation** | **% of total variation** | **P value** | **P value summary** | **Significant?** | |
| Interaction | 1.786 | 0.6222 | ns | No | |
| Row Factor | 0.3636 | 0.4439 | ns | No | |
| Column Factor | 83.23 | <0.0001 | **** | Yes | |
| **ANOVA table** | **SS (Type III)** | **DF** | **MS** | **F (DFn, DFd)** | **P value** |
| Interaction | 0.3996 | 10 | 0.03996 | F (10, 66) = 0.8075 | P=0.6222 |
| Row Factor | 0.08137 | 2 | 0.04068 | F (2, 66) = 0.8222 | P=0.4439 |
| Column Factor | 18.62 | 5 | 3.725 | F (5, 66) = 75.28 | P<0.0001 |
| Residual | 3.266 | 66 | 0.04948 | | |
| **Figure 4B** | | | | | |
| **ANOVA table** | **SS** | **DF** | **MS** | **F (DFn, DFd)** | **P value** |
| Treatment (between columns) | 0.3553 | 5 | 0.07105 | F (5, 18) = 18.52 | P<0.0001 |
| Residual (within columns) | 0.06907 | 18 | 0.003837 | | |
| Total | 0.4243 | 23 | | | |

| Figure 5A_Bj | | | | | |
|---|---|---|---|---|---|
| **Source of Variation** | **% of total variation** | **P value** | **P value summary** | **Significant?** | |
| Interaction | 15.49 | 0.0153 | * | Yes | |
| Row Factor | 1.694 | 0.2017 | ns | No | |
| Column Factor | 74.11 | <0.0001 | **** | Yes | |
| **ANOVA table** | **SS** | **DF** | **MS** | **F (DFn, DFd)** | **P value** |
| Interaction | 3523 | 10 | 352.3 | F (10, 18) = 3.204 | P=0.0153 |
| Row Factor | 385.4 | 2 | 192.7 | F (2, 18) = 1.752 | P=0.2017 |
| Column Factor | 16857 | 5 | 3371 | F (5, 18) = 30.66 | P<0.0001 |
| Residual | 1980 | 18 | 110 | | |

| Figure 5B_A431 | | | | | |
|---|---|---|---|---|---|
| **ANOVA table** | **SS** | **DF** | **MS** | **F (DFn, DFd)** | **P value** |
| Treatment (between columns) | 10783 | 5 | 2157 | F (5, 6) = 4.811 | P=0.0410 |
| Residual (within columns) | 2690 | 6 | 448.3 | | |
| Total | 13472 | 11 | | | |

| Figure S7_EGFR | | | | | |
|---|---|---|---|---|---|
| **Source of Variation** | **% of total variation** | **P value** | **P value summary** | **Significant?** | |
| Interaction | 20.35 | <0.0001 | **** | Yes | |
| Row Factor | 3.707 | 0.0039 | ** | Yes | |
| Column Factor | 51.29 | <0.0001 | **** | Yes | |
| **ANOVA table** | **SS** | **DF** | **MS** | **F (DFn, DFd)** | **P value** |
| Interaction | 5.736 | 5 | 1.147 | F (5, 60) = 9.908 | P<0.0001 |
| Row Factor | 1.045 | 1 | 1.045 | F (1, 60) = 9.022 | P=0.0039 |
| Column Factor | 14.45 | 5 | 2.89 | F (5, 60) = 24.96 | P<0.0001 |
| Residual | 6.947 | 60 | 0.1158 | | |

| Figure S7_pEGFR | | | | | |
|---|---|---|---|---|---|
| **Source of Variation** | **% of total variation** | **P value** | **P value summary** | **Significant?** | |
| Interaction | 0.968 | 0.8468 | ns | No | |
| Row Factor | 0.04965 | 0.75 | ns | No | |
| Column Factor | 71.87 | <0.0001 | **** | Yes | |
| **ANOVA table** | **SS (Type III)** | **DF** | **MS** | **F (DFn, DFd)** | **P value** |
| Interaction | 0.1029 | 5 | 0.02057 | F (5, 56) = 0.3999 | P=0.8468 |
| Row Factor | 0.005277 | 1 | 0.005277 | F (1, 56) = 0.1026 | P=0.7500 |
| Column Factor | 7.637 | 5 | 1.527 | F (5, 56) = 29.69 | P<0.0001 |
| Residual | 2.881 | 56 | 0.05144 | | |

| Figure S8_Bj | | | | | |
|---|---|---|---|---|---|
| **Source of Variation** | **% of total variation** | **P value** | **P value summary** | **Significant?** | |
| Interaction | 17.94 | 0.0005 | *** | Yes | |
| Row Factor | 11.91 | <0.0001 | **** | Yes | |
| Column Factor | 64.75 | <0.0001 | **** | Yes | |
| **ANOVA table** | **SS** | **DF** | **MS** | **F (DFn, DFd)** | **P value** |
| Interaction | 6438 | 10 | 643.8 | F (10, 18) = 5.982 | P=0.0005 |
| Row Factor | 4273 | 2 | 2136 | F (2, 18) = 19.85 | P<0.0001 |
| Column Factor | 23236 | 5 | 4647 | F (5, 18) = 43.18 | P<0.0001 |
| Residual | 1937 | 18 | 107.6 | | |

| Figure S8_SA24 | | | | | |
|---|---|---|---|---|---|
| **Source of Variation** | **% of total variation** | **P value** | **P value summary** | **Significant?** | |
| Interaction | 10.6 | 0.6407 | ns | No | |
| Row Factor | 4.055 | 0.248 | ns | No | |
| Column Factor | 61.15 | 0.0002 | *** | Yes | |
| **ANOVA table** | **SS** | **DF** | **MS** | **F (DFn, DFd)** | **P value** |
| Interaction | 2103 | 10 | 210.3 | F (10, 18) = 0.7881 | P=0.6407 |
| Row Factor | 804.8 | 2 | 402.4 | F (2, 18) = 1.508 | P=0.2480 |
| Column Factor | 12137 | 5 | 2427 | F (5, 18) = 9.097 | P=0.0002 |
| Residual | 4803 | 18 | 266.8 | | |

| Figure S8_SA48 | | | | | |
|---|---|---|---|---|---|
| **Source of Variation** | **% of total variation** | **P value** | **P value summary** | **Significant?** | |
| Interaction | 5.055 | 0.8368 | ns | No | |
| Row Factor | 2.164 | 0.3345 | ns | No | |
| Column Factor | 76.06 | <0.0001 | **** | Yes | |
| **ANOVA table** | **SS** | **DF** | **MS** | **F (DFn, DFd)** | **P value** |
| Interaction | 1464 | 10 | 146.4 | F (10, 18) = 0.5440 | P=0.8368 |
| Row Factor | 626.6 | 2 | 313.3 | F (2, 18) = 1.164 | P=0.3345 |
| Column Factor | 22023 | 5 | 4405 | F (5, 18) = 16.37 | P<0.0001 |
| Residual | 4843 | 18 | 269.1 | | |

**Supplementary data 1. Time-lapse video of the proliferation of A431 and Bj-5tα cells.** Treatment with 100 nM ligand was performed at the 24h mark. Evidence of a balance between differentiation and proliferation are found in the fibroblast, where a reduced proliferation is associated to cellular morphology differences in WT and K48T.
